# Targeting THOC2-Mediated mRNA Export Induces PARP Inhibitor Vulnerability in DNA Repair-Competent Hepatocellular Carcinoma

**DOI:** 10.64898/2026.05.17.725613

**Authors:** Xinli Li, Songpeng Yang, Mengxin Zhang, Zhaoyu Guo, Yan Wang, Yanru Meng, Yuanping Liu, Hu Zhang, Kaikun Xu, Xiuyuan Zhang, Yuanjun Zhai, Jingzhuo Jin, Fuchu He, Chunyan Tian, Aihua Sun

**Author notes:** Corresponding authors: Fuchu He; Chunyan Tian; Aihua Sun. These authors contributed equally (Xinli Li, Songpeng Yang, Mengxin Zhang).

## Abstract

Hepatocellular carcinoma (HCC) remains a lethal malignancy with limited therapeutic options. While Poly (ADP-ribose) polymerase inhibitors (PARPi) exploit synthetic lethality in tumors with DNA repair defects, their clinical utility in HCC is hindered by the low prevalence of canonical repair gene mutations and the enhancing DNA repair capacity. Through proteomic analysis of two independent cohorts (*n*=260), we identified the THO complex component THOC2 as a master regulator of DNA damage response (DDR) via mRNA nuclear export control. Clinically, THOC2 overexpression predicted poor survival (HR=2.68-6.84, *P*<0.001) and correlated with enhanced DDR gene expression. Mechanistically, THOC2 chaperones mRNA nuclear export of DDR effectors (MDC1, PRKDC, MSH6) and proliferation drivers (TOP2A), thereby establishing a dual pro-repair/pro-growth program. Targeting this vulnerability, THOC2 knockdown induced synthetic lethality with PARPi, reducing Olaparib IC50 by up to 61% and suppressing tumor growth by 76% (*P*<0.001). Our study illuminates mRNA transport as a druggable DDR modulator and establishes THOC2 as both a prognostic biomarker and a therapeutic target to overcome PARPi resistance in HCC. This work pioneers a strategy to expand synthetic lethality beyond genetic defects by targeting post-transcriptional regulation.

## Background

Hepatocellular carcinoma (HCC), responsible for over 700,000 annual deaths globally, exemplifies a lethal malignancy with dire therapeutic limitations[1]. Despite advancements in systemic therapies, first-line tyrosine kinase inhibitors (e.g., sorafenib, lenvatinib) yield marginal survival benefits (median OS extension: 3 months; ORR <6%) [2–6] , while PD-1-mediated immunotherapy only benefits 10-20% of patients[7, 8]. Critically, over 70% of patients relapse post-resection, underscoring the unmet need for novel therapeutic targets or combination therapies to improve the prognosis of HCC patients[9].

A paradigm-shifting approach in oncology exploits synthetic lethality by targeting DNA damage repair (DDR) vulnerabilities, exemplified by PARP inhibitors (PARPi) in BRCA-deficient tumors[10–12]. PARPs are enzymes activated by single- and double-strand breaks in DNA (SSBs and DSBs, respectively), catalyzing the transfer of ADP-ribose from nicotinamide adenine dinucleotide (NAD^+^) onto protein substrates[13, 14]. PARPi induces the accumulation of SSBs, leading to the collapse of replication forks during the S phase and subsequently to DSBs[15]. In HR-deficient tumors, such as BRCA1/2-deficient cells, these DSBs are predominantly repaired by error-prone non-homologous end joining (NHEJ), leading to excessive DNA damage and ultimately cell death[16]. PARPi also causes the trapping of PARP1 and PARP2 enzymes in damaged DNA, forming PARP–DNA complexes and ultimately generating severe cytotoxicity[17, 18]. Consequently, PARPi induces synergistic lethality in HR-deficient cells. However, HCC exhibits minimal HR deficiency (e.g., rare BRCA1/2 mutations), and its therapeutic resistance is primarily mediated by aberrant DDR protein overexpression[19]. Uncovering that targeting non-classical DDR hyperactivation drivers to induce a ’BRCAness’ phenotype may uncover novel therapeutic vulnerabilities and establish strategies to sensitize HCC to PARPi.

In this study, we identified THOC2—a master regulator of mRNA export—as a dual orchestrator of HCC progression and response to PARPi. Proteomic data from multiple independent cohorts revealed that THOC2 overexpression in majority of tumors, correlating with poor survival (HR=2.68-6.84, *P*<0.001). Mechanistically, THOC2 selectively exports mRNA encoding key DDR proteins (MDC1, PRKDC, MSH6), thereby sustaining HR proficiency. Crucially, knockdown of THOC2 sensitized HCC cells to PARPi. Our work positions THOC2 as a druggable linchpin connecting mRNA transport to DDR activation, offering a biomarker-driven strategy to overcome PARPi resistance in HCC.

## Methods

### Data availability

We collected three published HCC proteomics datasets, namely those by ‘Jiang et al.’s cohort’[20], ‘Gao et al.’s cohort’[21], and ‘Xing et al.’s cohort’[22]. The proteomic datasets from 14 distinct tumor types, including GBM[23], HNSCC[24], EC[25, 26], LUAD[27–30], LSCC[31], SCLC[32], BRCA[33], GC[34, 35], iCC[36], PDAC[37], CRC[38], ccRCC[39–41], HGSC[42, 43], are from Supplementary Information of corresponding papers. The TCGA whole-genome sequencing data for nine tumor cohorts, including HNSCC, ESCA, LUAD, LSCC, BRCA, LIHC, CRC, KIRC, and BLCA, was downloaded from the UCSC Xena website (https://xenabrowser.net/datapages/).

All data are available in the main text or the Supplementary material. The RIP sequencing data of THOC2 in Huh7 cells are available at the National Center for Biotechnology Information (PRJNA1119719). The Proteomic data of mouse subcutaneous tumors are available at the PRID (PXD052837, https://www.ebi.ac.uk/pride/, Username; Password: V9irPDMw74rg).

### Plasmids and reagents

The THOC2 overexpression plasmid (pCMV-MCS-3*Flag-THOC2) was obtained from Shandong Wei Zhen Biotechnology Co. Ltd (China). HisScript II 1st Strand cDNA Synthesis Kit (Cat#R211-02, Vazyme, China); Taq Pro Universal SYBR qPCR Master Mix (Cat#Q712-02, Vazyme); CCK-8 Cell Counting Kit (Cat#A311-02, Vazyme); BeyoClick™ EdU Cell Proliferation Kit with Alexa Fluor 555(Cat#C0075S, Beyotime, China); Fluorescent In Situ Hybridization Kit(Cat#C10910, RiboBio, China); FITC Annexin V Apoptosis Detection Kit I(Cat#556547, BD Pharmingen, USA); Olaparib (AZD2281, Selleck, USA). **Antibodies**

Primary antibodies against the following proteins were obtained from Proteintech (China): THOC1 (10920-1-AP, 1:500), THOC2 (55178-1-AP, 1:500, for WB), THOC5 (14862-1-AP, 1:500), THOC6 (15316-1-AP, 1:500), THOC7 (17881-1-AP, 1:500), TOP2A (20233-1-AP, 1:500), PRKDC (28534-1-AP, 1:500), MSH6 (18120-1-AP, 1:500), β-actin (66009-1-Ig, 1:3000), β-tubulin (66240-1-Ig, 1:3000); from Novus Biologicals (USA): MDC1 (NB100-395, 1:2000), THOC3 (NBP1-92502, 1:1000); from Merck Millipore (Germany): γH2AX (16-202A, 1:200 for IF); from Thermo Fisher: Goat anti-Rabbit IgG (H+L) Cross-Adsorbed Secondary Antibody, Alexa Fluor™ 488 (A-11008, 1:200).

### Difference Analysis

The Welch’s t-test was employed to identify differentially expressed proteins (DEPs) between groups. To account for multiple comparisons, *P*-values were adjusted using the False Discovery Rate (FDR) method. The log-fold change (logFC) is computed by taking the logarithm of the ratio between the means of each group.

For ‘Homology Directed Repair’ related functions, the sample population was divided into high and low expression groups based on the best cutoff value of ‘Homology Directed Repair’ ssGSEA (single-sample Gene Set Enrichment Analysis) score for OS calculated with survminer/ surv_cutpoint package in R. Significantly differentially expressed proteins between ‘Homology Directed Repair’ score high and low groups were calculated following the criteria: FDR < 0.05 and FC (high/low) > 1 in Jiang et al.’s cohort; FDR < 0.05 and FC (high/low) > 1 in Gao et al.’s cohort. For TREX related functions, the difference between the top 25% and bottom 25% of samples was analyzed based on the ssGSEA scores of TREX. Significantly differentially expressed proteins between TREX score high and low groups were calculated following the criteria: FDR < 0.05 and FC (high/low) > 2 in Jiang et al.’s cohort; FDR < 0.05 and FC (high/low) > 1.2 in Gao et al.’s cohort. Different FC thresholds were empirically selected according to the magnitude of proteomic differences within each comparison to ensure a sufficient number of differentially expressed proteins for robust downstream enrichment analyses while maintaining statistical rigor through FDR control. For screening differential proteins in the proteomic data of mouse subcutaneous tumors, the following criteria were utilized: *P* < 0.05 and logFC (shTHOC2/shCtrl) <-0.5.

### GO biological process and KEGG pathway enrichment analysis

Pathway enrichment analysis was performed using Metascape Gene Annotation & Analysis Resource[44] that uses several ontology sources: KEGG Pathway, GO Biological Processes, Reactome Gene Sets, Canonical Pathways and CORUM (https://metascape.org/gp/index.html#/main/step1) . The pathway with a significance of adjusted *P* value < 0.05 were defined as significantly regulated. GSEA analysis was performed in R/Bioconductor package fgsea (version1.24.0) or clusterProfiler (version 4.6.0) referring to the hallmark gene sets H, curated gene sets (C2) and ontology gene sets (C5) downloaded from the MSigDB database (http://www.gsea-msigdb.org/gsea/msigdb/index.jsp). Gene sets overlapping at least ten genes with the DEPs and with a significance of *P* < 0.05 were defined as significantly enriched. ssGSEA algorithm implemented in the R/Bioconductor package GSVA (version 1.46.0) was used to quantify the enrichment scores of the indicated gene sets for tumor and adjacent tissues. Heatmaps in this study were plotted using the R/Bioconductor package ComplexHeatmap (version 2.14.0). The enrichment network diagram is generated using the enrichmentNetwork function available in the aPEAR package[45] (version 1.0.0) in the R language. This function facilitates the creation of a network diagram based on enriched gene sets or pathways, providing a visual representation of the relationships between these biological processes.

### Weighted gene co-expression network analysis (WGCNA)

The R package ‘WGCNA’[46] was used for identifying key modules in Jiang and Gao et al.’s cohort. Differential proteins between tumor and adjacent tissues were identified for module detection using specific screening criteria (FDR < 0.01, LogFC > 1 in the Jiang et al.’s cohort; FDR < 0.01, LogFC > 0.2 in the Gao et al.’s cohort). The optimal soft threshold was determined based on the scale-free topology model fitting (R² = 0.8). The functional annotation of each module was performed using Metascape Gene Annotation & Analysis Resource.

### Correlation analysis

In the analysis, Spearman correlation coefficient was utilized to compute correlations between proteins. When *P*-value is less than 0.05, a correlation coefficient greater than 0 is interpreted as a positive correlation. Conversely, when *P*-value is less than 0.05, a correlation coefficient less than 0 is considered a negative correlation. Correlation heatmaps was generated using the corrplot package (Version 0.92) in the R language. This package provides functions to visualize correlation matrices in a heatmap format.

### Survival Analysis

Kaplan-Meier plots (Log-rank test) were used to describe OS or DFS in R software. The survival curves were calculated with the R function survfit from the R package survival (version 3.5-0) with the formula Surv (time, vitalstatus) ∼ categorie and plotted with the R function ggsurvplot from the R package survminer (version 0.4.9). The surv_cutpoint function from the survminer package is utilized to determine the optimal cut-off value. Subsequently, the power of the "coxph" function was harnessed to construct a Cox Regression model and compute Hazard Ratios (HR). HR > 1 and *P* < 0.05 indicate an increased risk of the event, HR < 1 and *P* <0.05 indicate a decreased risk, and a HR of 1 or *P* ≥ 0.05 suggests no difference in risk between groups.

### Cell culture

Huh7, MHCC97H, Hepa1-6 and HEK293T cells were obtained originally from the National Collection of Authenticated Cell Cultures. Huh7, MHCC97H and HEK293T maintained in Dulbecco’s modified Eagle’s medium (DEME, Gibco, USA) supplement with 10% fetal bovine serum (FBS, Newzerum, New Zealand) and 1% Pen/Strep (Gibco). Hepa1-6 was cultured in DMEM/F12 (Gibco) with 15% FBS (Gibco) and 1% Pen/Strep. All the cell lines were certificated by STR and free of mycoplasma contamination. All cells used for experiments were maintained in a humidified incubator at 37°C with 5% CO_2_ cells within 20 generations.

### Animal studies

Mice were maintained in the specific pathogen-free (SPF) facility of the National Center for Protein Sciences, Beijing, China (NCPSB) and housed under a 12h/12h light/dark cycle, with free access to regular chow and water. All mice were grouped by no more than 5 mice per cage. Animal use and care were approved by the NCPSB in accordance with the Institutional Animal Care and Use Committee Guidelines (IACUC-20220803-55MT). NOD-SCID and C57BL/6J mice were purchased from Charles River (China). All mice used in this study were male and aged between 4 and 5 weeks.

### Construction and transfection of shRNA and recombinant plasmids

The small interference RNAs (siRNAs) targeting THOC1, THOC2, THOC3, THOC5, THOC6, or THOC7 were purchased from Shanghai Genephama Co., Ltd. (China). The THOC2-targeting shRNAs were purchased from Tsingke Biotechnology Co., Ltd. (China). The lentiviral vector for stable knockdown of THOC2, based on the pLKO.1 backbone, was constructed using standard molecular cloning techniques. All primers used for plasmid construction are listed in the Oligonucleotides Table. For transfection, all siRNAs and THOC2-plasmids were transfected using Lipofectamine 3000 (Invitrogen, USA) according to the standard protocol. Briefly, in each 6-well plate, 2 μg of siRNA or plasmid was transfected. After transfection in serum-free medium for 12 h, the medium was replaced by serum-containing medium, and cells were harvested after an additional 36 h of culture. A highly efficient lentiviral system was used to generate the viruses: HEK-293T cells were transfected with target/scramble plasmids, psPAX2 packaging plasmid and pMD2.G envelope plasmid with PEI. After 48 h, lentivirus was harvested and transduced into target cells, which were then screened by 2 mg/ml puromycin (Gibco) for 3 d.

siTHOC1: 5’-GAUACCAAACCUACGAGAATT-3’

siTHOC2: 5’-GAGUUGUCAUAUCAUGUAATT-3’

siTHOC3: 5’-GUAAGAACCCUCAGUUUCATT-3’

siTHOC5: 5’-ACAUUGACGUUGUCCUGAATT-3’

siTHOC6: 5’-AGAAGCCGGUGGUGACUUUTT-3’

siTHOC7: 5’-GGAGAUGAUCGGAGAAUUATT-3’

THOC2-sh#1: 5’-GAGTTGTCATATCATGTAATT-3’

THOC2-sh#2: 5’-GGTCAGATCAGAAACACUATT-3’

### RNA extraction and RT-qPCR

Total RNA was extracted with Trizol reagent (Sigma, USA), and the RNA concentration was measured with Nanodrop2000 (Thermo Scientific, USA). First strand cDNA was reversed with HiScript II 1st Strand cDNA Synthesis Kit (Vazyme); and realtime polymerase chain reaction (PCR) were performed with Taq Pro Universal SYBR qPCR Master Mix (Vazyme) according to the procedure manual. GAPDH was used for mRNA normalization, and the primers used are listed in Oligonucleotides Table. The relative gene expression was calculated using the 2^−ΔΔCT^ method.

THOC1 Forward Primer 5’-GGAACACATGCCCACTTTGGAG-3’

THOC1 Reverse Primer 5’-AAGTGAGGGCTTCTCCGTGCTA-3’

THOC2 Forward Primer 5’-GGTAATCTTTCAGGAAGGTGGAGA-3’

THOC2 Reverse Primer 5’-GCTGATGTCATCCCAGACTTTG-3’

THOC3 Forward Primer 5’-TTCGCATCTGGGATGTGAGGAC-3’

THOC3 Reverse Primer 5’-GTTGCCTACAGCAATGGTCTGC-3’

THOC5 Forward Primer 5’-TCCAGAGGACTCCCAAGATGAC-3’

THOC5 Reverse Primer 5’-GTGCCTCTTCAGCATCTCCTTG-3’

THOC6 Forward Primer 5’-GTTTGGCAACTGATTCCGACTGG-3’

THOC6 Reverse Primer 5’-AGGTCCTGGTAGAAGGTGACGT-3’

THOC7 Forward Primer 5’-TGTAGCATAGCTGGAGCACATGA-3’

THOC7 Reverse Primer 5’-ATGCCTGTCTGGATGGTGCTGA-3’

GAPDH Forward Primer 5’-GTCTCCTCTGACTTCAACAGCG-3’

GAPDH Reverse Primer 5’-ACCACCCTGTTGCTGTAGCCAA-3’

β-actin Forward Primer 5’-AGAGCTACGAGCTGCCTGAC-3’

β-actin Reverse Primer 5’-AGCACTGTGTTGGCGTACAG-3’

TOP2A Forward Primer 5’-GAAGATGTTGGCATTAACTATAGGG-3’

TOP2A Reverse Primer 5’-CCGCAGCATTAACTAGAATCTC-3’

PRKDC Forward Primer 5’-CTTACATGCTAATGTATAAGGGCG-3’

PRKDC Reverse Primer 5’-CAGCAGGCACTTTACTTTCTC-3’

MDC1 Forward Primer 5’-TGCTCTTCACAGGAGTGGTG-3’

MDC1 Reverse Primer 5’-GGGCACACAGGAACTTGACT-3’

MSH6 Forward Primer 5’-GGCTCGAAAGACTGGACTTATT-3’

MSH6 Reverse Primer 5’-CCAGGAGGCTCTGTTCATTT-3’

### Western Blotting (WB)

Protein samples from HCC cells were extracted using RIPA lysis buffer (Solarbio, China) supplemented with 1× Halt protease and phosphatase inhibitor cocktail (Merck, USA). Equal amounts of protein (20–30 µg) were separated on 10% SDS-PAGE gels, transferred to 0.45 µm NC membrane (Pall Corporation, USA), and blocked with 5% BSA. Following an overnight incubation with primary antibodies (Abs) at 4 °C, the protein bands were further incubated with secondary antibodies. Unless otherwise stated, antibodies were diluted at 1:1000. Protein lanes were visualized using enhanced chemiluminescence substrate (NCM biotech, China) and exposed in the ImageQuant 800 system (Cytiva, USA). β-tubulin served as an internal control.

### Cell Proliferation Assay

Cells were seeded in a 96-well plate at a density of 3 × 10^3^ cells per well and cultured for 4 d. Cell proliferation was assessed using the Cell Counting Kit-8 (CCK8) assay (Beyotime). Briefly, 10 µL of CCK-8 solution was added to each well and incubated with the cells for 4 h at 37 °C in the dark. Subsequently, the absorbance of each well was measured at 450 nm using the Spark® multimode microplate reader (Tecan, Switzerland). Cell proliferation rate was presented as the fold change in absorbance.

### Colony Formation Assay and EdU Cell Proliferation Assay

Cells were seeded at a density of 3 × 10^3^ cells per well of a 6-well plate and culture them at 37°C for approximately 2 weeks. Following incubation, fix the cells with methanol for 20 min, stain with 0.5% crystal violet for another 20 min, wash with water twice, and allow to air-dry. Subsequently, photograph the plates and count the number of single colonies. Cell proliferation was assessed using the standard protocol provided with the EdU kit purchased from Beyotime Company. All assays were performed in three independent biological replicates, with each replicate consisting of at least three technical replicates. Data are presented as mean ± SEM.

### Subcutaneous transplantation and drug treatment

Male NOD-SCID mice at 5 to 6 weeks of age were injected subcutaneously on each flank with 5 × 10^6^ Huh7 suspended in 200 µL DMEM, while C57BL/6J mice received bilateral subcutaneous injections of 5 × 10^6^ Hepa1-6 cells suspended in 200 µL DMEM/F12. Tumor volume was assessed by measuring two diameters with digital calipers and calculated as x*x*y/2, where x is the smaller diameter and y is the larger diameter. For subcutaneous tumors in drug-treated mice, 5 × 10^6^ Hepa1-6 cells were injected into the subcutaneous right side of C57BL/6J mice. When tumors reached a volume of 200 mm^3^, mice were randomized to divided into four groups (6 mice each group) for treatment: (1) shCtrl treated with vehicle control (DMSO); (2) shCtrl treated with Olaparib (50 mg/kg); (3) shTHOC2 treated with vehicle control (DMSO) and (4) shTHOC2 treated with Olaparib (50 mg/kg). Tumor-bearing mice were injected intraperitoneal every day until sacrificed. Tumor size was monitored every other day, and experiments were terminated when the tumor volume had increased to 2000 mm^3^.

### Tissue Microarray (TMA) and Immunohistochemistry (IHC)

The HCC tissue microarrays were purchased from Shanghai Outdo Biotech (HLivH180Su11). And immunohistochemistry staining with anti-THOC2 (Abcam, 1:500) was performed by Shanghai Outdo Biotech (China). The TMA was evaluated and scored by a pathologist. Immunohistochemical staining of xenograft tumor tissues were performed by Wuhan Servicebio Technology with the following antibodies overnight: anti-ki67 (Servicebio, 1:600), anti-γH2AX (Merck, 1:200) and stained with goat anti-mouse IgG Ab (Servicebio, 1:200) at room temperature for 50 min and counterstained with hematoxylin.

### Immunofluorescence (IF)

1×10^6^ cells were plated on Glass Bottom Confocal Dishes (NEST, China), fixed with 4% paraformaldehyde (PFA) for 15 min at room temperature and permeabilized with 0.5% Triton X-100 for 15 min. The cells were incubated in blocking buffer 3% bovine serum albumin (BSA) in PBS for 50 min at room temperature and then with the indicated primary mouse monoclonal anti-γH2AX antibody (Millipore, 1:200) and secondary antibodies (Abcam, 1:200) at 4 °C overnight or at RT for 60 min, respectively. DAPI (Sigma) was used to stain the nucleus. The dishes were observed using a fluorescence microscope (Zeiss, Germany).

### Cell apoptosis assays

Huh7 or MHCC97H cells were treated with IR (6 Gy) for 12 h or Olaparib (0.8 µM) dissolved in DMSO for 48 h. The cell apoptosis was detected with an FITC Annexin V Apoptosis Detection Kit I according to the manufacturer’s instructions (BD Pharmingen, USA). Wash cells twice with cold PBS and then resuspend cells in 1X Binding Buffer at a concentration of 1×10^6^ cells/ml. Transfer 100 µl of the solution (1×10^5^ cells) to a 5 ml culture tube. Add 5 µl of FITC Annexin V and 5 µl PI. Gently vortex the cells and incubate for 15 min at RT (25°C) in the dark. Add 400 µl of 1×Binding Buffer to each tube and samples were analyzed with a flow cytometer (BD Accuri^TM^ C6, CA, USA). Each measurement was performed in triplicate and each experiment was carried out at least three times. Data are presented as mean ± SEM. **Alkaline Comet Assay**

Alkaline comet assays were performed to detect the genomic instability. Briefly, cells were harvested and resuspended in PBS at a concentration of approximately 1×10^5^ cells/ml. A small aliquot (10 μl) of the cell suspension was mixed with 120 μl of 0.5% low-melting-point agarose and immediately spread onto pre-coated microscope slides. The slides were then incubated at 4 °C in the dark for 10 minutes to allow the agarose to solidify. Next, the slides were immersed in lysis solution (2.5 M NaCl, 100 mM EDTA, 10 mM Tris, 1% Triton X-100, pH 10) and stored at 4 °C for 1 night to enable cell lysis and protein removal. Following lysis, the slides were placed in a horizontal electrophoresis tank filled with alkaline buffer (300 mM NaOH, 1 mM EDTA, pH > 13) for 20 minutes to unwind the DNA. Electrophoresis was then performed at 25 V and 300 mA for 20 minutes at 4°C. After electrophoresis, slides were neutralized with 0.4 M Tris buffer (pH 7.5) for 5 minutes and washed with distilled water. The slides were then stained with 1×SYBR Golid stain buffer (Thermo, USA) and visualized under a fluorescence microscope. Comet tail lengths and tail moments were measured as indicators of DNA damage using OpenComet in ImageJ (version 1.53k).

### RNA Immunoprecipitation Seq (RIP-seq)

The RIP experiment and RNA library construction and sequencing were conducted by Wuhan SeqHealth Tech Co., Ltd. (China). The THOC2 antibody (55178-1-AP) used in the experiment was purchased from Proteintech.

### Mouse Subcutaneous Tumors Proteomic Data Processing

5 paired samples of mouse subcutaneous tumors from the shCtrl and shTHOC2 groups were for mass spectrometry. Peptides (∼1 µg) obtained from tumors were used for LC-MS/MS analysis. The LC-MS/MS analysis was performed using an Q Exactive HF Hybrid Quadrupole-Orbitrap Mass Spectrometer (Thermo Fisher Scientific, USA). The MS system uses Data Independent Acquisition (DIA) for scanning, employing a time gradient of 150 min. The raw LC-MS/MS data files were analyzed using MaxQuant (version 2.1.2.0) with the spectra searched against the UniProt human database. Search parameters were set as follows: variable modifications: oxidation of methionine M, protein N-terminal acetylation; fixed modifications: Carbamidomethyl C; digestion mode: Trypsin/P; all other parameters were set to default values. After data quality control, 6 485 proteins from shCtrl samples and 6286 proteins from shTHOC2 samples were obtained and subjected to further differential analysis and functional enrichment analysis.

### RNA Fluorescence in Situ Hybridization/Immunofluorescence (FISH/IF)

RNA-FISH assay was performed with the Fluorescent in Situ Hybridization Kit (RiboBio). After fixation with 4% PFA, permeabilization was performed using 0.5% Triton-X100. The RNA probe (RiboBio) was added and incubated at 37°C overnight. Subsequently, the probe was washed with SSC of different concentrations, followed by blocking with 5% BSA. Next, the THOC2 antibody was added and incubated at 4°C overnight. After washing with 1× PBS, the fluorescent secondary antibody was applied and incubated at room temperature for 50 min. Finally, the samples were incubated with DAPI before imaging.

### IC₅₀ assay

HCC cells were seeded in 96-well plates at a density of 4,000 cells per well in 100 μL of culture medium and allowed to adhere overnight. Cells were then treated with serial dilutions of PARPi in culture medium. Each concentration was tested in three technical replicates. After 48 h of incubation at 37 °C with 5% CO₂, 10 μL of CCK-8 solution was added to each well, followed by incubation for an additional 2 h at 37 °C. Absorbance was measured at 450 nm with a reference wavelength of 650 nm using a microplate reader. Background absorbance from blank wells (medium plus CCK-8 without cells) was subtracted from all measurements. IC₅₀ values were calculated using nonlinear regression in GraphPad Prism. All experiments were performed at least three independent times. Data are presented as mean ± SEM.

### Statistical analysis

All experiments for quantitative analysis and representative images were reproduced with similar results for at least three times and the data were presented as mean ± SEM. Data sets with normal distribution were analyzed with student’s t test to compare continuous variables of 2 groups. Kaplan-Meier and log-rank test were constructed survival curves. All the statistical analyses were performed using R (version 4.2.1). and GraphPad Prism (version 9.5.1). *P* value of less than 0.05 were considered statistically significant (ns, *P* ≥ 0.05, *, *P* < 0.05; **, *P* < 0.01; ***, *P* < 0.001, ****, *P* < 0.0001).

## Results

### Proteomic data analysis reveals ‘Transport of Mature Transcript to Cytoplasm’ as a Potential Regulator of HR in HCC

To systematically evaluate the activation status of DNA repair pathways across cancers, we analyzed 23 published proteomics datasets spanning 14 types of tumors. Tumor-versus-adjacent tissue differential expression results followed by GSEA analysis revealed significant enrichment of DNA repair pathways—including mismatch repair (MMR), nucleotide excision repair (NER), base excision repair (BER), and double-strand break repair, particularly homologous recombination (HR) repair- in HCC (Figure 1A and Table S1). Consistent with this, ssGSEA scores for these pathways were markedly upregulated in tumors and strongly associated with poor prognosis (Figure S1A, B). Meanwhile, key DNA repair proteins were significantly upregulated in tumors compared to adjacent tumor tissues, particularly within the prognostically unfavorable S-II/III or S-Me/Pf subtype (Figure S1C). These proteomic signatures of enhanced DNA repair activity in HCC suggests a possible mechanistic basis for PARPi insensitivity, whereas tumors with intrinsic DNA repair defects, classically represented by PARPi-sensitive breast and ovarian cancers, demonstrate the expected inverse correlation between DNA repair activity and PARPi sensitivity (Figure 1A).

**Figure 1.**
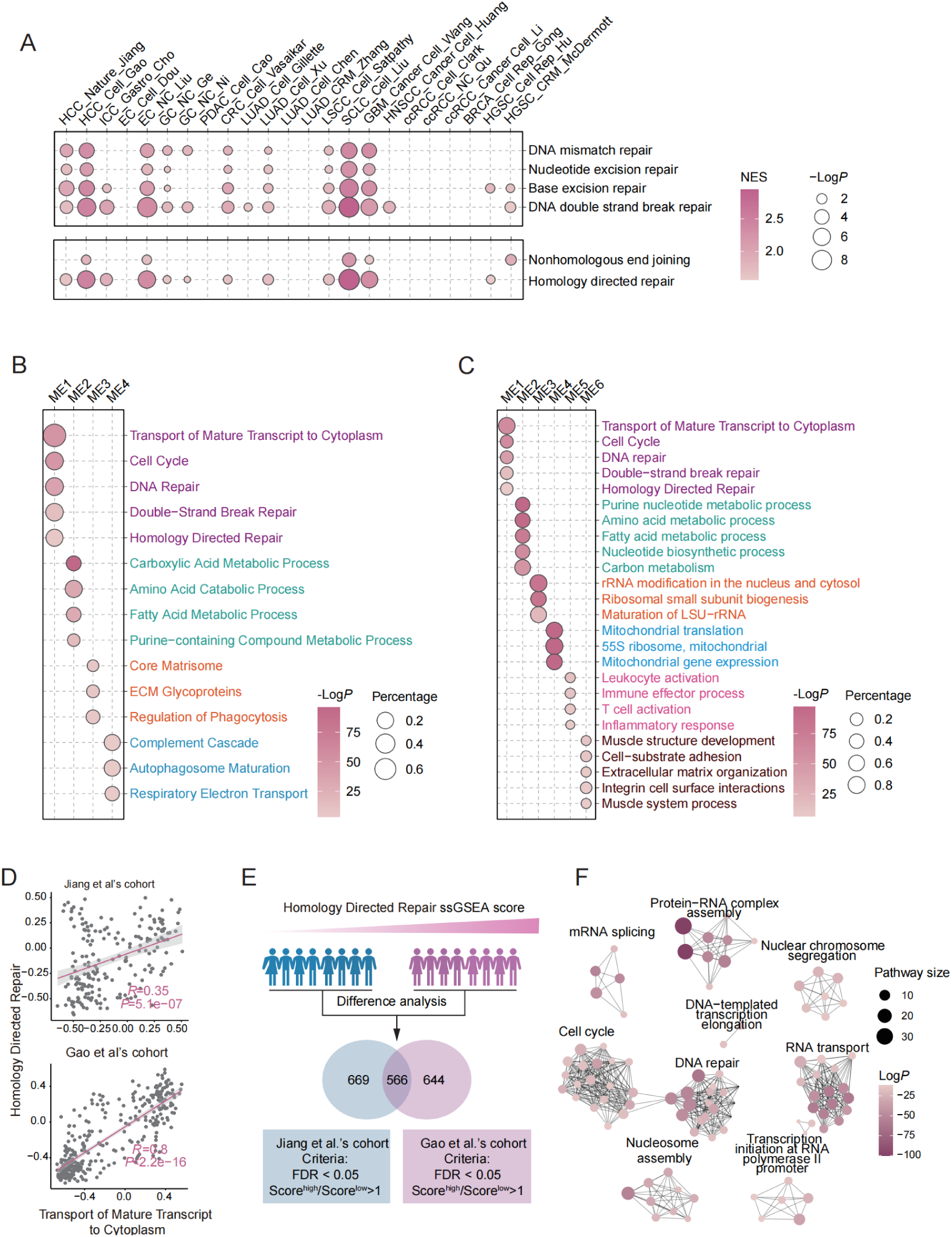
The ‘Transport of Mature Transcript to Cytoplasm’ regulates HR in HCC. **(A)** Pathway enrichment analysis of tumor-adjacent differential expression in DNA damage repair pathways using pan-cancer proteomics data. (**B, C)** Pathway enrichment analysis of different modules identified in Weighted Gene Co-Expression Network Analysis (WGCNA) from the Jiang (**B)** and Gao et al.’s cohorts (**C)**. (**D)** Scatter plot showing the correlation between ssGSEA scores of the ‘Homology Directed Repair’(HDR) pathway and the ‘Transport of Mature Transcript to Cytoplasm’ pathway. (**E)** Schematic of the analytical workflow for identifying HDR-active HCC subgroups. The population was stratified into high and low HDR groups based on ssGSEA scores, followed by differential protein expression analysis. A Venn diagram illustrates overlapping proteins from the Jiang and Gao et al.’s cohorts, with screening criteria detailed below. (**F)** Pathway enrichment analysis of the proteins jointly screened (**E)**. Circle size represents the number of proteins contained per pathway, and color intensity indicates the enriched *P*-value. Permutation test for (**A)**; Fisher’s exact test for **(B, C** and **F)**; Spearman correlation coefficient for (**D)**; two-tailed unpaired Welch test for (**E)**.

To further investigate the regulatory mechanisms of DNA repair in HCC, we conducted Weighted Gene Co-Expression Network Analysis (WGCNA) on two independent proteomic datasets to identify co-expression modules associated with DNA repair. Strikingly, the ‘Transport of Mature Transcript to Cytoplasm’ was significantly enriched within the DNA repair module, exhibiting the most robust statistical significance among all enriched pathways (Figure 1B, C and Table S1). To further validate this association, we calculated the correlation between ‘Transport of Mature Transcript to Cytoplasm’ and the DNA repair related pathway using ssGSEA scores. Intriguingly, both proteomic datasets revealed a strong positive correlation, suggesting a close association between these pathways (Figure 1D and Figure S1D, E). Notably, similar positive correlations between RNA export-related signatures and DNA repair pathways were also observed in additional tumor types with highly activated DDR pathways, including esophageal carcinoma (EC), small cell lung cancer (SCLC), and glioblastoma (GBM), suggesting that this association may represent a broader regulatory feature across malignancies rather than being restricted to HCC (Figure S1F).

Next, we examined the proteomic characteristics of HCC patients with active HR repair and notably found that RNA export pathway-related proteins were significantly upregulated in this group (Figure 1E, F and Table S1). These findings, consistent with the WGCNA results, suggest a strong correlation between HR repair and RNA export. Clinically, the ‘Transport of Mature Transcript to Cytoplasm’ was only notably elevated in tumor tissues but also strongly correlated with unfavorable patient outcomes (Figure S1G, H).

In summary, our results demonstrate that DNA repair pathways, particularly HR repair, are highly active in HCC and indicate that a potential regulatory role for RNA export processes in modulating DNA repair efficiency.

### THOC2 Links Nuclear mRNA Export to DNA Repair in HCC

To further elucidate the functional relationship between ’Transport of Mature Transcript to Cytoplasm’ and DNA repair in HCC, we systematically classified proteins involved in the Transport of Mature Transcript to Cytoplasm pathway into three functional groups: those participating in mRNA biogenesis, primarily including mRNA elongation and poly-adenylation factors; the TREX complex responsible for nuclear transport of mRNA; and the nuclear pore complex (NPC) mediating cytoplasmic export. Strikingly, GSEA analysis comparing HR-active and HR-inactive populations, identifed the TREX complex as the only significantly enriched pathway (Figure 2A). In parallel analyses, the TREX complex emerged as the singularly enriched pathway in tumor tissues while simultaneously exhibiting the most pronounced upregulation relative to other export factors (Figure S2A, B). Further analysis revealed that beyond its canonical role in mRNA export, high TREX complex activity was associated with significant enrichment of DNA repair and cell cycle pathways, suggesting its broader functional involvement in HCC progression (Figure S2C, D and Table S2).

**Figure 2.**
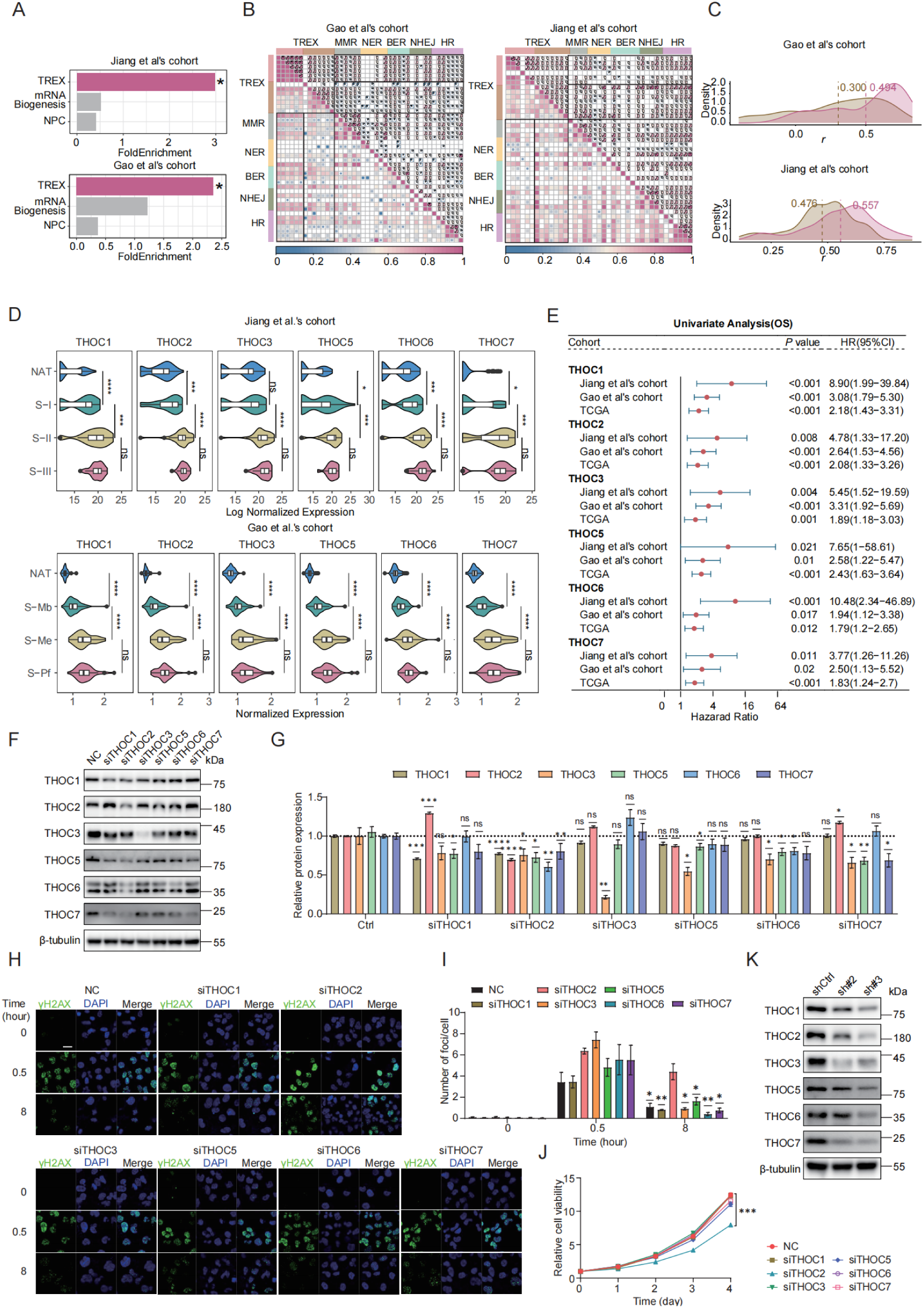
THOC2 regulates DNA damage repair in HCC. **(A)** Gene set enrichment analysis (GSEA) of mRNA biosynthesis, TREX, and nuclear pore complex (NPC) pathways in high versus low HR score groups. (**B)** The heatmap depicting the correlation between TREX proteins and DNA damage repair (DDR) proteins. (**C)** The density plot illustrates the distribution of correlation coefficients between the DNA repair proteins and the two components of TREX, and values represent the median correlation coefficient. Rose red represents THO complex, and yellow-brown represents other parts of TREX. (**D)** The violin plot displaying the expression of the THO complex in the two datasets. (**E)** The forest plot of survival analysis (OS) of THO complex in two proteomics data sets and TCGA. HR > 1 indicates worse prognosis with higher expression. (**F)** Western blot (WB) analysis of the knockdown effect of siRNA on THO complex proteins in Huh7 cells. **(G)** Densitometric quantification of THO complex protein expression levels shown in **(F)**. Protein expression levels were normalized to β-tubulin and presented relative to the control group (NC) (mean ± SEM; *n* = 3). (**H, I)** Immunofluorescence staining (IF) assay shows γH2AX foci in the control and THO complex knockdown cells after ionizing radiation (IR; 0.6 Gy, **H**). And histograms presented the average number of foci per cell in each group (mean ± SEM; *n* = 3, **I**). Scale bar, 20 μm. (**J)** CCK-8 assays were applied to evaluate proliferation abilities of Huh7 cells after transiently knockdown of the components of THO complex with their relative siRNA, respectively (mean ± SEM; *n* = 3). (**K)** WB analysis of THO complex expression in control and THOC2 stable knockdown Huh7 cells. Cellular β-tubulin expression served as the loading control. shCtrl: non-targeting shRNA; sh#2: sh-THOC2-2; sh#3: sh-THOC2-3; NC: negative control. Fisher’s exact test for (**A)**; Spearman correlation coefficient for (**B)**; two-tailed unpaired Welch test for (**D, G** and **I)**; Log-rank test for (**E)**; two-way ANOVA for **(J)**; ns no significance; \**P* < 0.05; ** *P* < 0.01; *** *P* < 0.001 and **** *P* < 0.0001.

Subsequently, correlation analysis revealed that among TREX subcomplexes, the THO complex exhibited the strongest internal consistency and associations with DNA repair proteins compared to other subcomplexes (Figure 2B, C). These findings identify the THO complex as the pivotal component of TREX complex mediating DNA repair functions. In addition, we observed that cell cycle-related pathways were significantly co-expressed with DNA repair and mRNA nuclear export pathways (Figure 1B, C). Consistent with this, correlation analysis between TREX proteins and proliferation markers identified the THO subcomplex as showing the most significant associations (Figure S2E), confirming its dual role in both DNA repair and cell cycle regulation within TREX. Moreover, the THO complex was consistently upregulated in tumors within the proteomics dataset and was strongly associated with shorter overall survival (OS; Figure 2D, E). Most THO components were also significantly correlated with shorter disease-free survival (DFS), with the exception of THOC5 in the Jiang et al.’s cohort and THOC7 in the Gao et al.’s cohort (Figure S2G). To further assess the robustness of these observations, survival analyses across multiple cutoff values (10th–90th percentile) were performed for THOC5 and THOC7 in both cohorts. Although statistical significance was not reached at some cutoff settings, high expression consistently showed a trend toward poorer prognosis, with HRs generally remaining >1 (Figure S3). A similar trend was observed in the TCGA data, further supporting the notion that the THO complex is markedly dysregulated in HCC (Figure S2F, G).

To further delineate the functional hierarchy within the THO complex, we used siRNA to knock down individual subunits (Figure 2F, G and Figure S2H) and evaluated their impact on DNA damage repair capacity. Strikingly, the knockdown of THOC2 resulted in the most substantial accumulation of γH2AX foci (Figure 2H, I), indicating severe DNA repair impairment. Additionally, CCK8 assays further revealed that THOC2 silencing exerted the strongest suppression of HCC cell proliferation (Figure 2J). As a known scaffold protein of the THO complex, THOC2 knockdown led to coordinated downregulation of other THO components (Figure 2K). Furthermore, poly(A)+ RNA FISH analysis demonstrated that depletion of THOC2 caused the most pronounced nuclear accumulation of poly(A)+ RNA among all THO components tested, indicating a dominant role in maintaining mRNA export activity (Figure S2I, J). In summary, these results establish THOC2 as the structural and functional linchpin of TREX complex, contributing to DNA repair and proliferative capacity in HCC.

### THOC2 as an independent prognostic predictor for HCC

We next assessed the potential of THOC2 as a prognostic marker for HCC. Notably, high THOC2 expression levels were strongly correlated with shorter OS and DFS in HCC (Figure 3A, B and Figure S4A, B). This consistent trend was also observed in Xing et al.’s cohort (Figure S4C, D). Furthermore, IHC results from an independent HCC cohort revealed that THOC2 expression was significantly elevated in tumors and correlated with poor prognosis (Figure 3C-F). The forest plot illustrates the hazard ratios (HR) for THOC2 levels in HCC patients across various clinical covariates (including AFP, MVI, and recurrence risk post-resection), suggesting that THOC2 provides a more accurate prognostic prediction than existing clinical indicators (Figure 3G and Figure S4E). We also analyzed the correlation between THOC2 and clinical indicators associated with malignancy, such as tumor diameter, tumor number, MVI, AFP, and BCLC stage. Surprisingly, THOC2 showed a significant correlation only with AFP in both datasets (Figure 3H and Figure S4F). This suggests that THOC2 may represent distinct malignant features.

**Figure 3.**
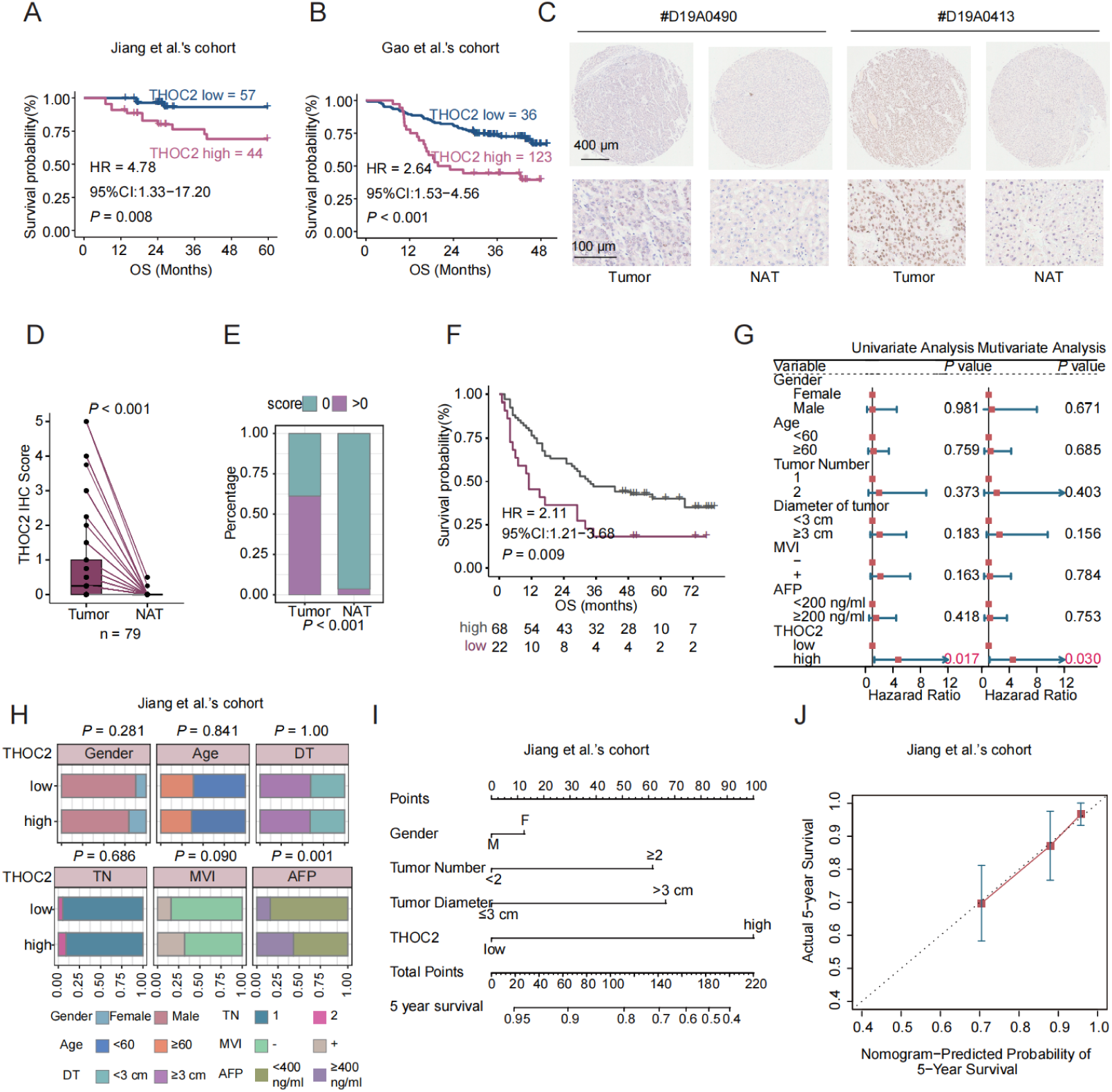
THOC2 as an independent prognostic predictor in HCC. (A, **B)** Kaplan-Meier overall survival (OS) curves stratified by THOC2 expression (optimal cut-off) in Jiang (**A)** and Gao et al.’s cohort (**B**. **C)** Representative immunohistochemistry (IHC) images of THOC2 in tumor and adjacent non-tumorous tissues (NAT). On the left: Samples with longer OS (57 months). On the right: Samples with shorter OS (6 months). Scale bars: 400 μm (top); 100μm (bottom). (**D, E)** THOC2 IHC scores were significantly higher in tumor than NAT (*n* = 79). (**F)** Kaplan-Meier analysis of the OS based on THOC2 IHC scores. Patients were stratified into low (score = 0) and high (score > 0). (**G)** Univariate and multivariate Cox regression analyses of THOC2 and clinical characteristics in the Jiang et al.’s cohort. (**H)** The correlation between THOC2 expression and clinical indicators in Jiang et al.’s cohort. Patients were stratified into high- and low-THOC2 groups based on the cut-off value of THOC2 expression in (**A)**. DT: diameter of tumor; TN: tumor number; MVI: microvascular invasion. (**I)** Nomogram constructed by combining THOC2 and clinical risk factors. (**J)** Calibration plots of the nomogram for predicting the probability of 5-year survival. Log-rank test for (**A, B** and **F)**; Two-tailed unpaired Wilcoxon t test for (**D)**; chi-square test for (**E, H)**.

To further evaluate the prognostic generalizability of THOC2 across different clinical contexts, subgroup survival analyses were performed in the Jiang and Gao et al.’s cohorts. In the Jiang et al.’s cohort, THOC2-high patients consistently exhibited poorer survival trends across subgroups stratified by AFP level, MVI status, and tumor size (Figure S4G). Significant prognostic differences were observed in the AFP <400 µg/L subgroup and in patients with tumor size ≥3 cm, while several other subgroups showed similar trends but did not reach statistical significance, likely due to limited sample sizes. Consistently, subgroup analyses in the Gao et al.’s cohort based on AFP level, tumor number, BCLC stage, and tumor size demonstrated a generally unfavorable prognosis associated with high THOC2 expression across most clinical strata (Figure S4H).

To further assess the robustness of THOC2 as a prognostic marker, survival analyses were additionally performed using multiple cutoff values ranging from the 10th to the 90th percentile of THOC2 expression. In the Gao et al.’s cohort, high THOC2 expression was associated with poorer overall survival across most cutoff settings, although the median cutoff did not reach statistical significance. In the Jiang et al.’s cohort, statistical significance was observed at selected cutoff values, including the median cutoff, while several other cutoffs showed similar survival trends without reaching significance, likely due to limited sample size and patient distribution. Importantly, hazard ratios remained consistently >1 across most cutoff settings in both cohorts (Figure S5), supporting the overall stability of the prognostic effect of THOC2.

Furthermore, combining THOC2 with clinical indicators improves the predictive accuracy for HCC prognosis (Figure 3I, J). Taken together, these findings underscore the potential of THOC2 as a valuable prognostic marker in HCC.

### THOC2 promotes HCC proliferation and DNA repair

Subsequently, we further confirmed the effect of THOC2 on HCC proliferation. We established the stable THOC2-knockdown cell lines in Huh7 and MHCC97H cells via lentivirus infection (Figure S6A, B). CCK8 assay indicated that the cell viability was significantly reduced after THOC2 knockdown (Figure 4A). Additionally, the number of clones and EdU-positive cells were significantly reduced in the knockdown group (Figure 4B-E). Conversely, overexpression of THOC2 led to a significant increase in cell proliferation and the number of EdU-positive cells (Figure S6C-E). Furthermore, re-expression of THOC2 in THOC2-knockdown cells effectively rescued the impaired proliferative capacity induced by THOC2 depletion (Figure S6F). Consistent with *in vitro* results, THOC2 knockdown could significantly reduce the subcutaneous tumor growth (Figure 4F-J and Figure S6G-J). These results suggested the crucial role of THOC2 in promoting HCC proliferation in *vivo* and *vitro*.

**Figure 4.**
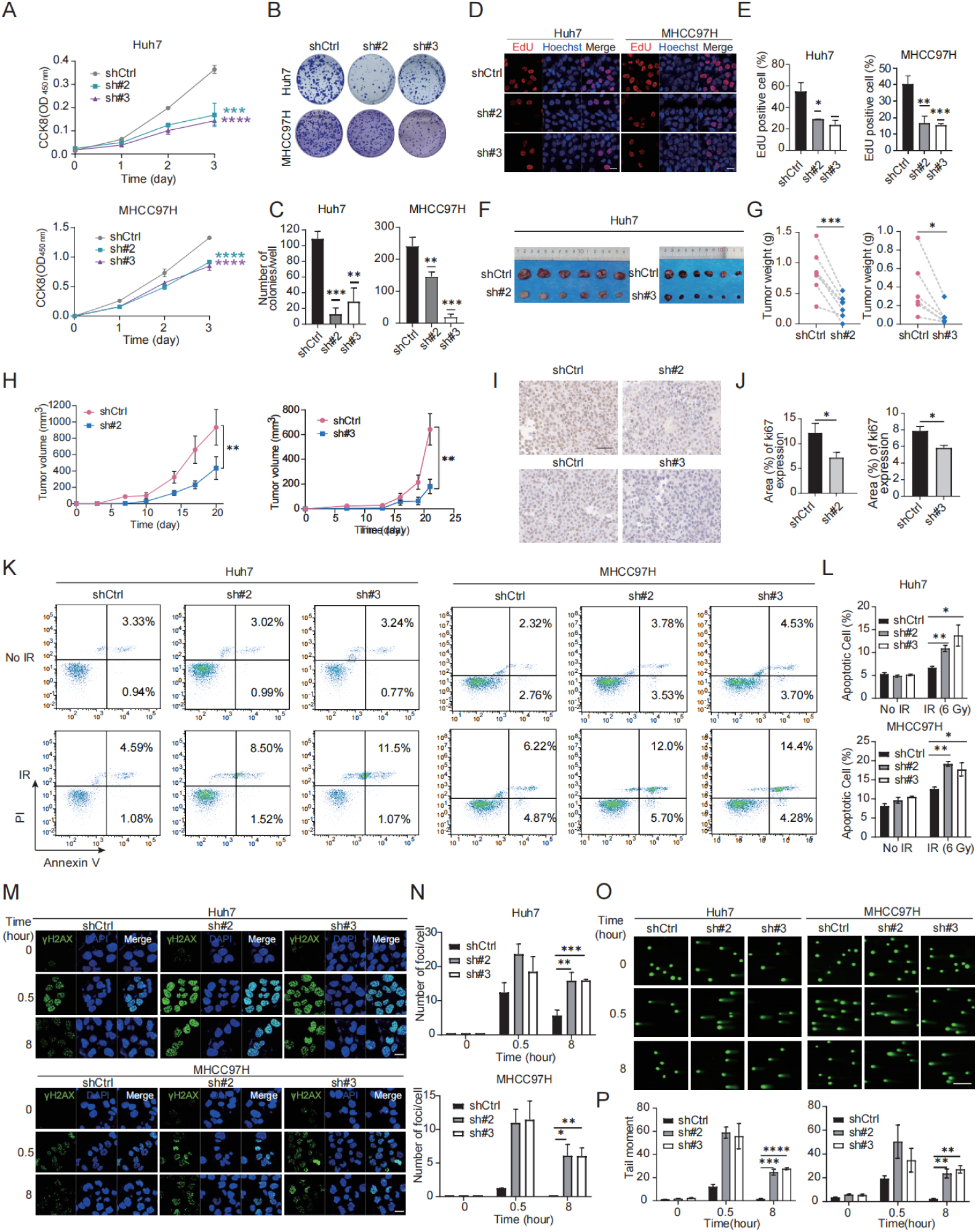
THOC2 promotes HCC proliferation and DNA repair. (A-E) CCK8 A, colony formation **(B, C)** and EdU assays **(D, E)** were applied to evaluate the proliferation abilities of control or THOC2-knockdown Huh7 and MHCC97H cells (mean ± SEM; *n* = 3). Scale bar, 50 μm. (**F-J)** Tumor xenograft model was constructed with stable control and THOC2-knockdown Huh7 cells (mean ± SEM; *n* = 6). Tumors were collected from sacrificed mice and tumor weights were measured (**G)**. Tumor sizes were recorded consecutively to establish tumor growth curves (**H)**. IHC staining of Ki67 in mouse subcutaneous tumors (**I-J)**. Scale bar, 50 μm. (**K, L)** Apoptosis experiment in the control and THOC2-knockdown cells after irradiation (IR) treatment (dose = 6 Gy). And histograms presented the percentage of apoptosis cells in each group (mean ± SEM; *n* = 3). (**M, N)** IF assay shows γH2AX foci in the control and THOC2-knockdown cells after IR treatment (dose = 6 Gy). And histograms presented the average number of foci per cell in each group (mean ± SEM; *n* = 3). Scale bar, 20 μm. (**O, P)** The alkaline comet assays show that the DSB repair capability is reduced in THOC2-knockdown Huh7 and MHCC97H cells. HCC cells are harvested at the indicated time (NT, and 0.5 h and 8 h post-treatment) upon IR treatment (6 Gy). The tail moment was analyzed using the CometScore software (mean ± SEM; *n* = 3). Scale bar, 200 μm. shCtrl: nontargeting shRNA; sh#2: sh-THOC2-2; sh#3: sh-THOC2-3; two-tailed unpaired Wilcoxon t test for (**A, C, E, L, N** and **P)**; two-tailed paired Wilcoxon t test for (**G, H** and **J)**; \**P* < 0.05; ** *P* < 0.01; *** *P* < 0.001 and **** *P* < 0.0001.

We also explored the effect of THOC2 on HCC DNA damage repair (DDR) activity. When cells were treated with irradiation (IR), THOC2 could decrease the cell apoptosis, suggesting that THOC2 plays a role in response to DNA damage (Figure 4K, L). Moreover, we observed that in response to IR, THOC2 knockdown significantly delays the clearance of γ-H2AX foci in Huh7 and MHCC97H cells (Figure 4M, N). The alkaline comet assays also revealed a substantial DNA repair defect in THOC2-knockdown Huh7 and MHCC97H cells at 8 h after treatment with IR (Figure 4O, P). Importantly, reintroduction of THOC2 markedly reduced γ-H2AX accumulation in THOC2-knockdown cells, indicating restoration of DNA repair capacity (Figure S6J, K). Together, these results suggest that THOC2 is a positive regulator of DDR.

### THOC2 facilitates mRNA export of the proliferation and DDR associated molecules

Considering the key function of THO complex in mRNA export, to explore the underlying mechanism by how THOC2 regulates proliferation and DDR, we established screening strategy to identify the target mRNAs of THOC2 for export. Firstly, we calculated the co-expressed proteins of THOC2 using the two proteomic datasets. 862 proteins demonstrated significant positive correlations with THOC2 in both datasets (Figure S7A and Table S3). Pathway enrichment analysis unveiled significant enrichment associated with mRNA splicing, cell proliferation, and DDR (Figure S7B and Table S3). Subsequently, we conducted RNA immunoprecipitation sequencing (RIP-seq) in Huh7 cells to identify the mRNA transcripts that interact with THOC2. The results showed that THOC2 predominantly binds to the coding sequence (CDS) region and 5’ untranslated region (UTR) of mRNA (Figure S7F). Remarkably, a total of 5,814 RNAs were significantly enriched by THOC2 antibodies (*P* < 0.05; Figure S7G and Table S3), with their functions primarily concentrated in crucial cellular processes such as chromatin organization, DNA repair, and regulation of cell cycle process (Figure S7H and Table S3). Additionally, we employed proteomic analysis on the aforementioned subcutaneous tumors (Figure 4F) with significantly reduced THOC2 expression (Figure S7C). A total of 596 proteins were identified as significantly downregulated in the THOC2-knockdown group (Figure S7D and Table S3), primarily involved in RNA metabolism, lipid metabolism, ribosomes and cell cycle, and DNA damage-related processes (Figure S7E and Table S3). A total of 111 targets were identified by intersecting the four data sets mentioned above (Figure 5A and Table S3). Besides RNA-related functions, these targets are primarily involved in DDR and cell cycle, aligning with the enrichment results from the different datasets mentioned above (Figure 5B and Table S3). This suggests that THOC2 may contribute to cell cycle and DNA repair in HCC by regulating the nuclear export of mRNA. Specifically, the proliferation marker TOP2A, pivotal for cell proliferation[47], along with the proteins MDC1[48, 49] and PRKDC[50, 51] involved in DNA double-strand break repair, and the key protein MSH6[52, 53] implicated in mismatch repair, were all among the identified proteins (Figure 5C, D). These mRNAs are significantly enriched by THOC2 (Figure 5E), and the expression of the corresponding proteins is markedly reduced after THOC2-knockdown (Figure 5F). Additionally, in the population cohort, these proteins show a significant positive correlation with the expression of THOC2 (Figure 5G). In conclusion, the results suggest that THOC2 may contribute to the progression of HCC by regulating the nuclear export of proliferation- and DNA repair-related mRNAs, such as TOP2A, MDC1, PRKDC, and MSH6.

**Figure 5.**
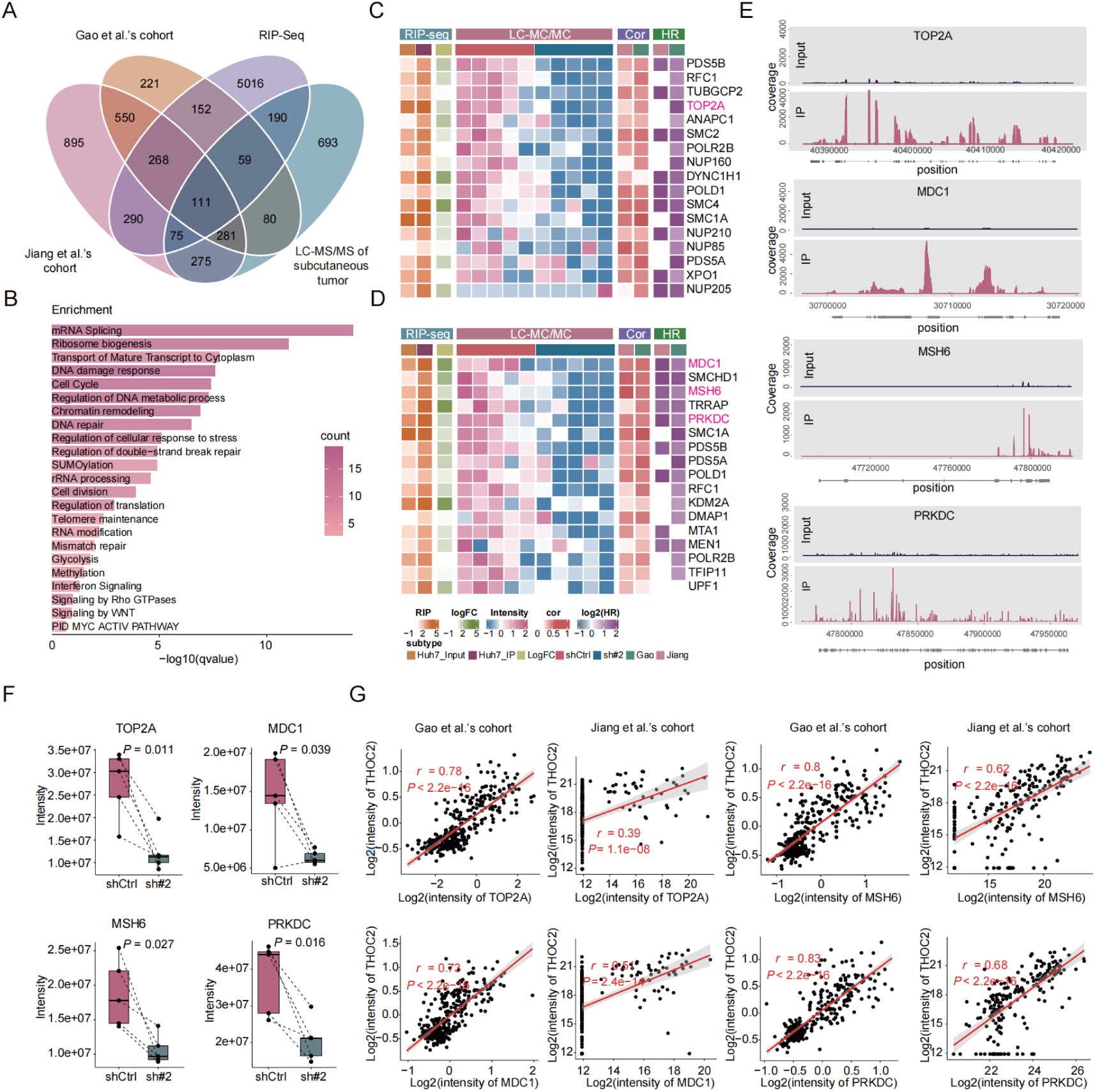
THOC2 selectively exports mRNAs related to proliferation and DNA repair. **(A)** Venn diagram for screening THOC2-selective export of mRNAs, wherein Jiang et al.’s cohort and Gao et al.’s cohort represent co-expressed proteins of THOC2 (*P* < 0.01; *r* > 0), RIP-Seq represents mRNA bound by THOC2 in the Huh7 cells (*P* < 0.05; IP/Input > 2), and LC-MS/MS of mouse subcutaneous tumors represents proteins down-regulated in the THOC2 knockdown group compared to the control group (in at least four samples of paired shTHOC2 and shCtrl samples, shTHOC2/shCtrl is less than 1). (**B)** Pathways enriched by intersection proteins in (**A)**. Color represents the number of intersecting proteins in pathways. (**C, D)** Heatmaps of proteins in cell cycle **(C)** and DNA repair **(D)** contained in intersection proteins. LC-MS/MS represents mass spectrometry data of mouse subcutaneous tumors, Cor represents correlation analysis with THOC2 and HR represents the relationship between protein and prognosis in the proteomic datasets. (**E)** RIP-Seq results of THOC2 binding to the selected mRNA in Huh7 cells. (**F)** Expression intensity of TOP2A, MDC1, MSH6, and PRKDC in proteomic data of mouse subcutaneous tumors (mean ± SEM; *n* = 5). (**G)** Correlation between TOP2A, MDC1, MSH6, PRKDC and THOC2 expression in Gao et al.’s cohort and Jiang et al.’s cohort, respectively. shCtrl: nontargeting shRNA; sh#2: shTHOC2-2; two-tailed paired Wilcoxon t test for (**F)**; Pearson correlation coefficient for (**G)**.

### THOC2 governs protein expression through nuclear export regulation of TOP2A, MDC1, PRKDC and MSH6 mRNA

To further confirm the regulatory effect of THOC2 on TOP2A, MDC1, PRKDC, and MSH6, we evaluated the protein expression levels in the THOC2 knockdown cell line. Our results showed that THOC2 knockdown significantly reduced protein levels of four proteins, whereas THOC2 overexpression significantly increased them (Figure 6A, B). However, whilst the total mRNA expression of TOP2A, MDC1, PRKDC, and MSH6 was not perturbed by THOC2 silence (Figure 6C, D). Additionally, we employed Fluorescence in Situ Hybridization/Immunofluorescence (FISH/IF) experiments to assess the regulatory impact of THOC2 on mRNA export from the nucleus. Within the nucleus, the co-localization of THOC2 and those four mRNAs indicated their interaction. At the same time, the fluorescence intensity of those four mRNAs in the cytoplasm was significantly reduced in the THOC2-knockdown cells compared to the control group (Figure 6E, F). This observation suggested that knockdown of THOC2 indeed inhibited the nuclear export of TOP2A, MDC1, PRKDC, and MSH6 mRNAs. In addition, nucleocytoplasmic fractionation followed by qRT–PCR analysis revealed a marked increase in the nuclear-to-cytoplasmic (N/C) ratio of all four transcripts upon THOC2 knockdown (Figure 6G), further supporting that THOC2 is required for efficient nuclear export of these mRNAs.

**Figure 6.**
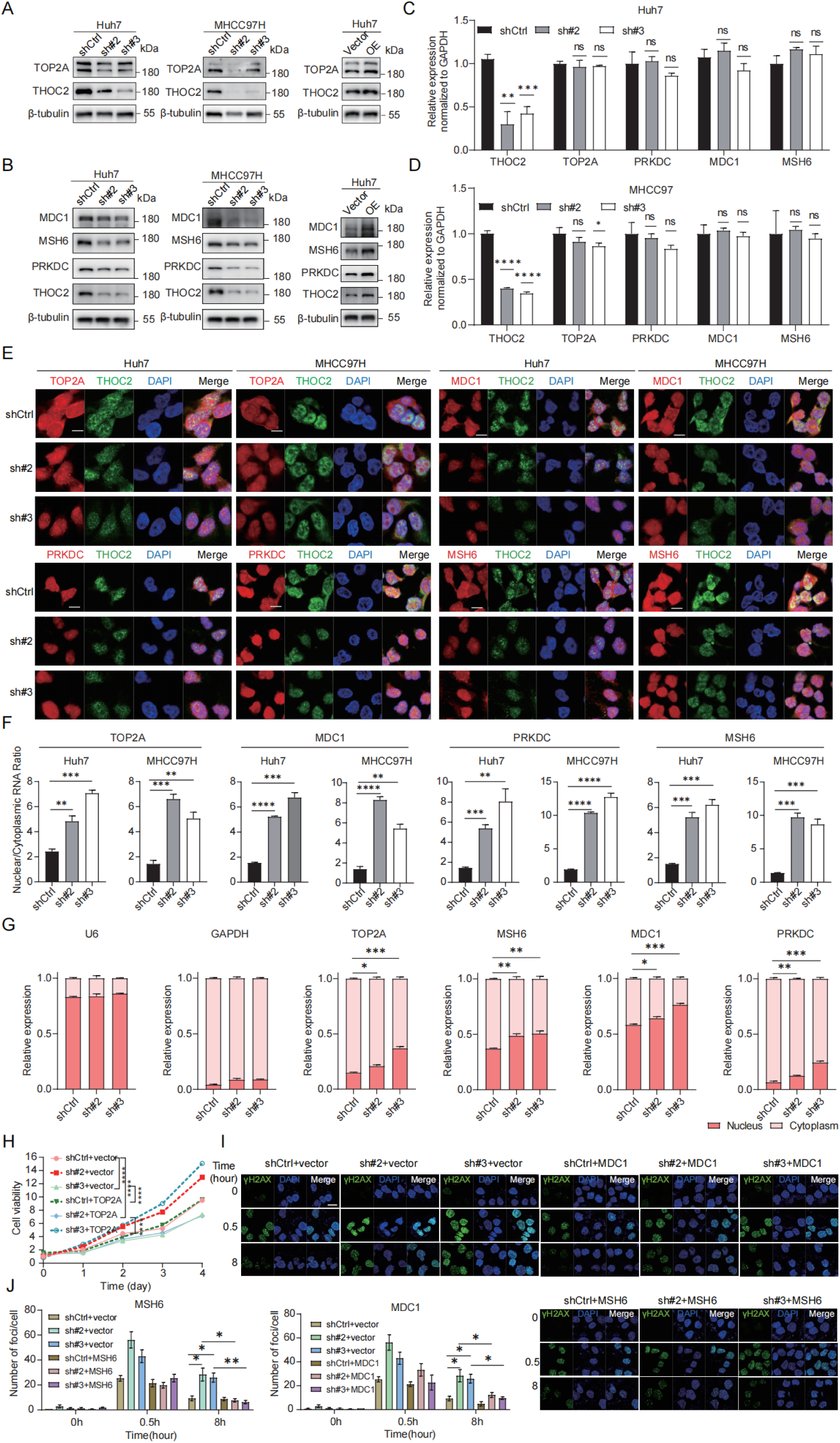
THOC2 facilitates the nuclear export of TOP2A, MDC1, PRKDC and MSH6 mRNA. (A, **B)** Protein level of TOP2A, MDC1, MSH6 and PRKDC was measured in THOC2-knockdown Huh7 and MHCC97H cells or THOC2-overexpressing Huh7 cells. (**C, D)** The total mRNA expression (mean ± SEM; *n* = 3) of TOP2A, PRKDC, MDC1, and MSH6 was detected by qRT-PCR in Huh7 **(C)** and MHCC97H cells **(D)**. (**E)** FISH/IF assays conducted on control and THOC2 knockdown cells of Huh7 and MHCC97H. Scale bar, 10 μm. **(F)** Quantification of nuclear/cytoplasmic RNA fluorescence intensity ratio in control and THOC2-knockdown Huh7 and MHCC97H cells (mean ± SEM, *n* = 3) based on FISH/IF assays shown in (E). **(G)** Nucleocytoplasmic fractionation followed by qRT–PCR analysis of THOC2 target mRNAs. Data are presented as nuclear-to-cytoplasmic (N/C) ratios (mean ± SEM, *n* = 3). **(H)** Cell proliferation assays in THOC2-knockdown MHCC97H cells following TOP2A overexpression (mean ± SEM, *n* = 3). **(I)** Representative immunofluorescence images of γH2AX foci in THOC2-knockdown MHCC97H cells with or without MDC1 or MSH6 overexpression. Scale bar, 20 μm. **(J)** Quantification of γH2AX foci per cell in (I) (mean ± SEM, *n* = 3). shCtrl: nontargeting shRNA; sh#2: shTHOC2-2; sh#3: shTHOC2-3; Vector: empty vector; OE: THOC2 overexpression; two-tailed unpaired Wilcoxon t test for (**C, D, F, G and J)**; two-way ANOVA for **(H)**; ns, no significance; \**P* < 0.05; ** *P* < 0.01; *** *P* < 0.001 and **** *P* < 0.0001.

To further validate the functional significance of THOC2 downstream targets, rescue experiments were subsequently performed in THOC2-knockdown HCC cells. Re-expression of TOP2A partially restored the proliferative capacity of THOC2-knockdown cells (Figure 6H), indicating that TOP2A contributes to THOC2-mediated promotion of HCC cell growth. In parallel, overexpression of the DNA damage response factors MDC1 or MSH6 markedly reduced γH2AX foci accumulation induced by THOC2 knockdown, as demonstrated by immunofluorescence staining and quantitative analysis of γH2AX foci per cell (Figure 6I, J). These rescue experiments validate that THOC2 exerts its functions at least in part through TOP2A, MDC1, and MSH6.

In summary, our investigation reveals that THOC2 exerts regulatory control over the expression of proteins associated with cell proliferation and DNA repair by modulating mRNA nuclear export.

### THOC2 knockdown induces HCC response to PARPi

In our study, THOC2 depletion might hinder HR, NHEJ, and MMR mechanisms by compromising the expression of key proteins such as MDC1, PRKDC, and MSH6, creating conditions conducive for PARP inhibitor (PARPi) treatment. In proteomic datasets, we observed a significant positive correlation between THOC2 and PARP1/2 (Figure S8A-D). Consistent with THOC2, both proteins were significantly upregulated in the more severe S-II/III or S-Me/Pf subtypes and were significantly associated with poor prognosis (Figure S8E-P). Therefore, we speculated that THOC2 depletion may synergize with PARPi, and we examined the half-maximal inhibitory concentration (IC50) of the approved PARPi Olaparib in the THOC2 knockdown Huh7, MHCC97H and Hepa1-6 cells. Indeed, we observed a decrease in the IC50 value of PARPi in the THOC2-knockdown group compared to the control group (Figure 7A-C). Moreover, knockdown of THOC2 enhanced PARPi-induced apoptosis (Figure 7D). To further evaluate the synergistic effect of THOC2 depletion and PARP inhibitor on DNA damage accumulation, we performed γH2AX immunofluorescence analysis *in vitro*. Notably, combined treatment with THOC2 knockdown and Olaparib resulted in a marked increase in γH2AX foci compared with either treatment alone, indicating enhanced DNA damage accumulation and impaired DNA repair capacity (Figure 7E, F).

**Figure 7.**
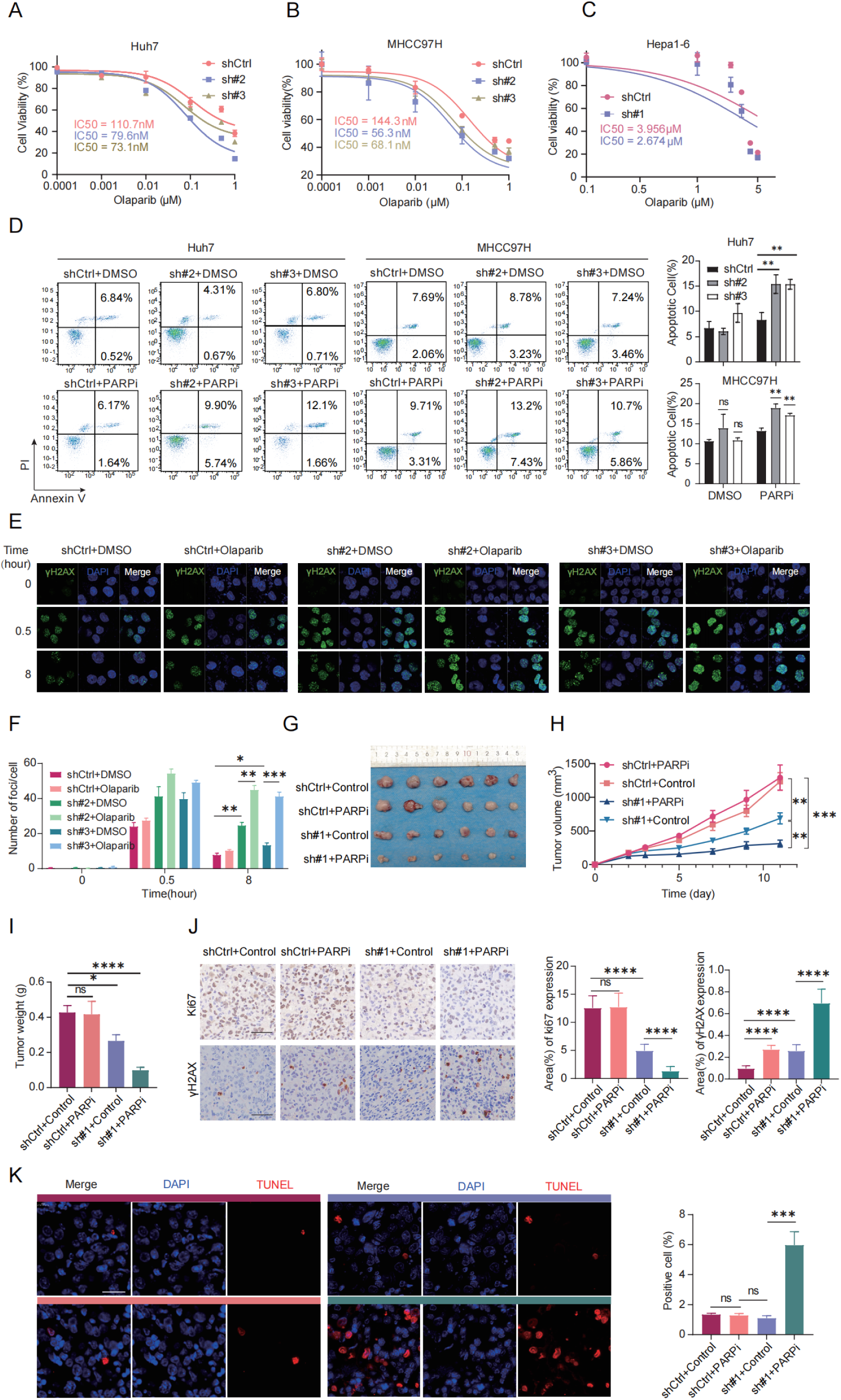
Targeting THOC2 induces HCC sensitivity to PARP inhibitors. (A-C) The dose-response curves of PARP inhibitor Olaparib on Huh7 **(A)**, MHCC97H **(B)** and Hepa1-6 **(C)** cells, both control and THOC2 knockdown, were generated using CCK8 assays (mean ± SEM; *n* = 3). The half maximal inhibitory concentrations (IC50) were shown. (**D)** Apoptosis experiment after THOC2 knockdown combined with PARPi (0.8 µM) in Huh7 and MHCC97H. PARPi: cells treated with Olaparib which is dissolved in DMSO; DMSO: cells treated with an equal volume of DMSO to Olaparib. **(E, F)** Immunofluorescence staining (IF) assay shows γH2AX foci in control and THOC2-knockdown MHCC97 cells treated with or without PARP inhibitor after ionizing radiation (IR; 0.6 Gy). And histograms presented the average number of foci per cell in each group (mean ± SEM; *n* = 3). Scale bar, 20 μm. (**G-I)** Subcutaneous Hepa1-6 cells tumors, tumor growth curves and tumor weight in the indicated treatment groups (mean ± SEM; *n* = 6). The dose of Olaparib used is 50 mg/kg. (**J)** Representative images of IHC Ki67 and γH2AX staining and the relative IHC scores (mean ± SEM; *n* = 6) in Hepa1-6 tumor tissues in (**G)**. Scale bar, 50 μm. (**K)** Representative images of TUNEL staining and the positive cells (%) (mean ± SEM; *n* = 6) in Hepa1-6 tumor tissues in **e**. Scale bar, 50 μm. The colors above the image represent the groups. shCtrl: nontargeting shRNA; sh#2: shTHOC2-2; sh#3: shTHOC2-3; sh#1: sh*Thoc2*-1; PARPi: mouses treated with Olaparib (50 mg/kg); Control: mouses treated with an equal volume of DMSO to Olaparib; two-tailed unpaired Wilcoxon *t* test for (**D, F, I, J** and **K)**; two-way ANOVA for **(H)**; ns, no significance; \**P* < 0.05; ** *P* < 0.01; *** *P* < 0.001 and **** *P* < 0.0001.

**Figure 8.**
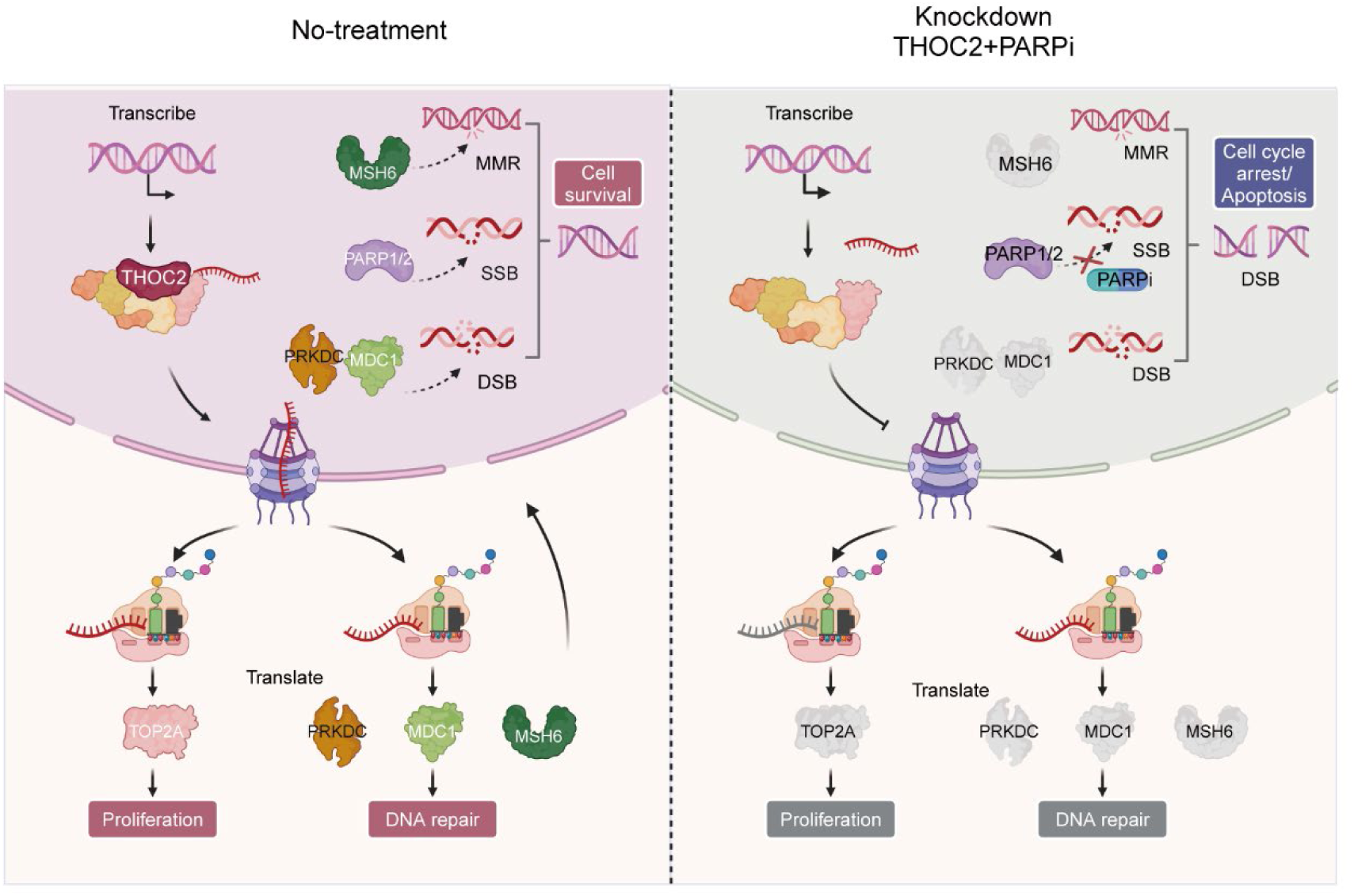
Model of the function and underlying mechanism of THOC2 in promoting HCC proliferation and DNA repair and sensitizing PARPi. The export of mRNA from the nucleus to the cytoplasm is a key step in protein synthesis and is essential for all eukaryotic cells. THO complex is crucial in the formation of co-transcriptional messenger ribonucleic acid particles that are transported to the cytoplasm for translation. THOC2, as the scaffold of THO complex, plays a key role in mRNA nuclear export. It regulates the nuclear export of mRNAs for key proteins involved in cell proliferation and DNA repair, such as TOP2A, MDC1, PRKDC, and MSH6, thereby increasing the expression of these proteins and promoting the progression of HCC. Targeting THOC2 in combination with PARP inhibitors can inhibit HCC proliferation and promote the occurrence of DSBs, inducing cell death. This combination represents a potential treatment for HCC patients with poor prognosis. (This figure was created with Biorender.com)

Importantly, reintroduction of THOC2 into THOC2-knockdown cells markedly attenuated the increased apoptotic response induced by PARPi treatment, further confirming that the enhanced PARPi sensitivity is specifically dependent on THOC2 loss rather than off-target effects (Figure S8R, S). To further assess whether this sensitizing effect extends beyond a single PARP inhibitor, we additionally evaluated the response to Niraparib and the DNA-damaging chemotherapeutic agent doxorubicin. THOC2 knockdown significantly enhanced the sensitivity of HCC cells to both Niraparib and doxorubicin, as reflected by reduced IC50 values (Figure S9A-B). Moreover, combined treatment with THOC2 knockdown and either Niraparib or doxorubicin markedly increased apoptotic responses compared with either treatment alone (Figure S9C-D). These findings suggest a potential enhanced sensitivity to PARPi treatment in cells with THOC2 depletion, highlighting the therapeutic implications of targeting THOC2 in HCC.

To further elucidate the synergistic role of THOC2 combined with PARPi in suppressing HCC *in vivo*, we initiated by utilizing THOC2-knockdown Hepa1-6 cells to establish a xenograft tumor model. Subsequently, intraperitoneal injections of DMSO and Olaparib were administered to the control group and THOC2-knockdown group, respectively. Notably, the combined treatment exhibited a more pronounced effect in inducing DNA damage and inhibiting tumor growth compared to single treatment alone (Figure 7E-G, Figure S10). In the combination group, tumor weight was reduced, accompanied by a decrease in Ki67-positive cells, indicative of decreased proliferation (Figure 7H). Conversely, γH2AX-positive cells and apoptotic cells, were significantly increased in the combination group (Figure 7H-I and Figure S8Q), underscoring the enhanced efficacy of combined therapy.

Taken together, our findings underscore the potential of THOC2 as a promising therapeutic target for HCC. Significantly, results from both *in vivo* and *in vitro* experiments demonstrate that inhibition of THOC2 synergizes effectively with PARPi, highlighting a promising therapeutic strategy for HCC.

## Discussion

The therapeutic potential of PARPi in HCC has been limited by the lack of canonical HR defects, such as BRCA1/2 mutations. However, recent studies have identified novel regulators of HR proficiency in HCC, offering new avenues to expand PARPi utility. For instance, NRDE2, a susceptibility gene identified through rare-variant association studies, promotes HR repair by enhancing casein kinase 2 (CK2) activity, which phosphorylates MDC1 to facilitate HR. NRDE2 deficiency sensitizes HCC cells to PARPi, particularly when combined with chemotherapy, highlighting its potential as a therapeutic target[54]. Similarly, the histone methyltransferase KMT5C has been shown to promote HR by facilitating RAD51/RAD54 complex formation, and its inhibition sensitizes HCC cells to PARPi[55]. These findings underscore the importance of targeting non-canonical HR regulators to overcome PARPi resistance in HCC.

In this context, our study identifies THOC2, a core mRNA export protein, as a multi-faceted regulator of HCC progression and PARPi sensitivity. Unlike NRDE2 and KMT5C, which primarily modulate HR through direct interactions with DDR proteins, THOC2 exerts broader oncogenic effects by selectively exporting mRNAs encoding not only DDR effectors (e.g., MDC1, PRKDC) but also mismatch repair factors (e.g., MSH6) and proliferation drivers (e.g., TOP2A, SMC3, POLD1). This unique dual role positions THOC2 as a central hub coordinating both genomic stability and tumor growth.

THOC2’s ability to regulate a wide array of oncogenic mRNAs sets it apart from other HR regulators. For example, TOP2A, an isoform of topoisomerase II (TOP2), is a nuclear protein crucial for DNA replication and cell division[47, 56–58]. Its overexpression is associated with aggressive tumor behavior and chemoresistance, making it a critical target in cancer therapy. Similarly, SMC3, PDS5 and SMC4, which are involved in chromosome cohesion and segregation, play pivotal roles in maintaining genomic integrity during cell division[59–62]. By exporting these mRNAs, THOC2 ensures the proper expression of proteins critical for tumor proliferation and survival.

In addition to proliferation-related proteins, THOC2 also regulates key DDR components. MDC1, a crucial mediator of the DNA damage response, recognizes double-strand break sites and recruits repair proteins to maintain genomic stability[63]. PRKDC, another THOC2 cargo, is essential for NHEJ, a major DSB repair pathway[64]. Additionally, MSH6, a component of the MMR sites system, ensures the fidelity of DNA replication by correcting base-pairing errors[65]. By controlling the expression of these DDR proteins, THOC2 maintains the genomic integrity necessary for tumor survival under stress conditions. Importantly, rescue experiments demonstrated that re-expression of MDC1 or MSH6 partially alleviated the DNA damage accumulation induced by THOC2 knockdown, further supporting the notion that THOC2 contributes to multiple DNA repair pathways through regulation of key repair factors. These findings suggest that THOC2-mediated mRNA export may coordinately influence HR, NHEJ, and MMR-related processes rather than acting on a single DDR pathway. Nevertheless, although our data support a broad role of THOC2 in DNA repair regulation, the precise contribution of THOC2 to individual repair pathways remains incompletely resolved. Due to technical limitations, we were unable to establish stable and reproducible HR/NHEJ reporter systems in the current study, and PRKDC rescue experiments could not be completed because of challenges associated with its large coding sequence. Future studies incorporating pathway-specific functional assays and structural analyses will therefore be necessary to further dissect the mechanistic role of THOC2 in distinct DDR pathways.

Furthermore, RIP-Seq data revealed that THOC2 also binds to mRNAs encoding key proteins involved in HR, such as BRCA1, BRCA2, and PALB2[66]. Although these proteins were excluded from the initial screening due to their expression levels falling below the detection limit of proteomics, their interaction with THOC2 suggests a potential role in fine-tuning HR activity. This finding aligns with the broader function of THOC2 as a master regulator of mRNA export, capable of modulating multiple DDR pathways simultaneously.

Compared to NRDE2 and KMT5C, which primarily target specific DDR components, THOC2’s ability to regulate a diverse set of oncogenic mRNAs provides a more comprehensive therapeutic target. For instance, while NRDE2 enhances HR by promoting MDC1 phosphorylation and KMT5C facilitates RAD51/RAD54 complex formation, THOC2 coordinates the expression of multiple DDR and proliferation-related proteins, offering a broader impact on tumor biology. This multifaceted regulatory capacity suggests that targeting THOC2 could yield more robust therapeutic benefits, including enhanced PARPi sensitivity and direct suppression of tumor growth.

Beyond its role in HCC, THOC2 is known to regulate vital biological processes, including the differentiation of vascular smooth muscle cells [67] and embryonic stem cells[68], through the selective mRNA export. However, its prognostic significance and mechanistic contributions in solid tumors, particularly in HCC, remain incompletely understood. Our study addresses this gap by demonstrating that THOC2 overexpression correlates with poor prognosis and functions as an independent prognostic predictor in HCC. Additionally, we reveal that THOC2’s oncogenic effects are mediated through its selective export of mRNAs encoding key DDR and proliferation-related proteins, providing a mechanistic basis for its role in HCC progression.

These findings suggest a central role of THOC2 in maintaining THO complex stability and function. However, we acknowledge an important limitation that the precise structural hierarchy and interdependence among THO complex components cannot be fully resolved in the present study. In addition, due to the lack of tools enabling selective disruption of complex assembly, we cannot completely exclude partial redundancy among subunits or potential THO complex-independent functions of THOC2. Further structural and mechanistic studies will be required to address these aspects.

The synthetic lethality between THOC2 deficiency and PARPi underscores the therapeutic promise of targeting mRNA export in HCC. However, the absence of THOC2-specific inhibitors remains a critical barrier. Future efforts should prioritize high-throughput drug screening and structure-based inhibitor design to translate THOC2’s mechanistic insights into clinical applications. Additionally, validating THOC2 as a predictive biomarker for PARPi response in prospective clinical trials could pave the way for precision therapy in HCC.

In summary, our study underscored the significant dysregulation of the mRNA nuclear export pathway in the development and progression of HCC, with THOC2 emerging as a pivotal player in this process. Notably, THOC2 stood out as a potential prognostic marker for HCC, and targeting it sensitized HCC cells to PARP inhibitors. Our findings provide novel insights into the molecular mechanisms driving HCC development and progression, offering potential new therapeutic avenues for HCC patients. THOC2 emerges as a promising therapeutic target, and further research into its targeting may yield innovative treatment strategies for HCC.

## Abbreviations

HCC: hepatocellular carcinoma
PARPi Poly (ADP-ribose): polymerase inhibitors;
OS: overall survival
DFS: disease-free survival
ssGSEA: single-sample gene set enrichment analysis
HNSC: Head-Neck Squamous Cell Carcinoma
ESCA: Esophageal carcinoma
BRCA: Breast carcinoma
LUAD: Lung adenocarcinoma
LSCC: Lung squamous cell carcinoma
CRC: Colorectal cancer
BLCA: Bladder Urothelial Carcinoma
NAT: normal tissues adjacent to the tumor
TREX: transcription-export
NPC: the nuclear pore complex
FC: foldchange
THO: complex suppressors of the transcriptional defects of hpr1Δ by overexpression
THOC2: THO Complex Subunit 2
MVI: microvascular invasion
AFP: alpha-fetoprotein
CCK-8: cell counting kit-8
WB: western blot
IHC: immunohistochemistry;
DDR: DNA damage repair
IR: Irradiation
γH2AX: histone H2AX phosphorylation at serine 139
RIP-seq: RNA immunoprecipitation sequencing
MDC1: Mediator of DNA Damage Checkpoint 1
PRKDC: Protein Kinase, DNA-Activated, Catalytic Subunit
MSH6: MutS Homolog 6
FISH/IF: fluorescence in situ hybridization/immunofluorescence
HR: homologous recombination
NHEJ: non-homologous end joining
MMR: mismatch repair
IC50: the half-maximal inhibitory concentration
DSB: DNA double-strand break

## Acknowledgements

We thank the National Center for Protein Sciences (Beijing) Animal Platform, the FCM platform, the imaging platform, and the MS platform of the for their assistance with mouse tumor models, FCM analysis, microscopy imaging, and MS analysis.

## Authors’ Contributions

A.S., C.T., and X.L. designed the study. A.S., C.T., and F.H. supervised the project. X.L. performed bioinformatic analyses. X.L. performed major experiments with assistance from S.Y. and M.Z.. Z.G., Y.M., H.Z., Y.L., X.Z. and Y.Z. provided technical and/or material support. X.L. and M.Z. wrote the manuscript. A.S. and C.T. revised the manuscript.

## Ethics approval and consent to participate

The animal care and experimental protocols were approved by the Institutional Animal Care and Use Committee (IACUC) of National Center for Protein Sciences (Beijing), Ethical review number: IACUC-20220803-55MT.

## Availability of data and materials

All data are available in the main text or the Supplementary material. The RIP sequencing data of THOC2 in Huh7 cells are available at the National Center for Bio-technology Information (PRJNA1119719). The Proteomic data of mouse subcutaneous tumors are available at the PRID (PXD052837, https://www.ebi.ac.uk/pride/, Username; Password: V9irPDMw74rg).

## Consent for publication

All authors have read and approved the final version of this manuscript.

