## Supplementary material for "Targeting THOC2-Mediated mRNA Export Induces PARP Inhibitor Vulnerability in DNA Repair-Competent Hepatocellular Carcinoma": Revised_Supporting_Information.docx

**Figures 1-6**

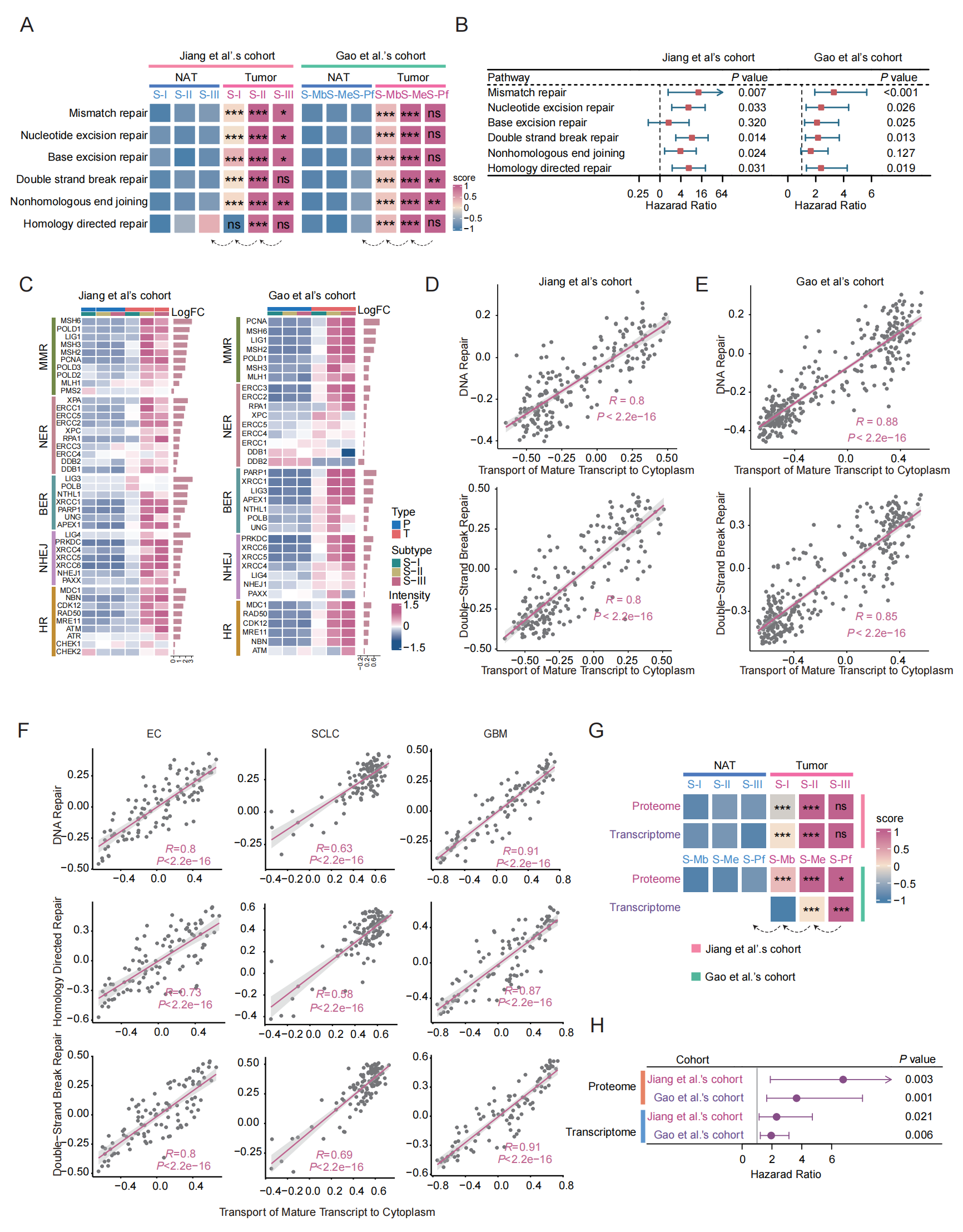

**Figure S1.** **Transport of mature transcript to cytoplasm plays a role in the DNA damage repair process.** (**A)** The heatmap depicting ssGSEA scores of DNA repair related pathways in NAT (normal tissue adjacent to the tumor) and tumor subtypes within proteome of Jiang et al.’s cohort and Gao et al.’s cohort. The color of each cell represents the average ssGSEA enrichment score; red denotes activation and blue denotes suppression. The arrow below indicates that the *P*-value of this column is calculated with respect to the column pointed to by the arrow. (**B)** Forest plot of survival analysis of ssGSEA score in (**A)**. HR > 1 indicates that a higher score is associated with a worse prognosis. (**C)** Heatmap illustrating the expression of DNA repair proteins in tumor and NAT samples. The histogram on the right displays the log fold change (logFC) between tumor and NAT samples. (**D, E)** Scatter plot showing the correlation between ssGSEA scores of DNA repair related pathways and the ‘Transport of Mature Transcript to Cytoplasm’ pathway in Jiang et al.’s cohort **(D)** and Gao et al.’s cohort **(E).** **(F)** Correlation between RNA export-related pathways and DNA repair programs across esophageal carcinoma (EC), small cell lung cancer (SCLC) and glioblastoma (GBM). **(G)** The heatmap depicting ssGSEA scores of ‘Transport of Mature Transcript to Cytoplasm’ in NAT and tumor subtypes within proteome and transcriptome of Jiang et al.’s cohort and Gao et al.’s cohort. The color of each cell represents the average ssGSEA enrichment score; red denotes activation and blue denotes suppression. The arrow below indicates that the *P*-value of this column is calculated with respect to the column pointed to by the arrow. **(H)** Forest plot of survival analysis of ssGSEA score in **(G)**. HR > 1 indicates that a higher score is associated with a worse prognosis. Two-tailed unpaired Welch test for (**A** and **G)**; Log-rank test for (**B** and **H)**; Spearman correlation coefficient for (**D**, **E** and **F)**.

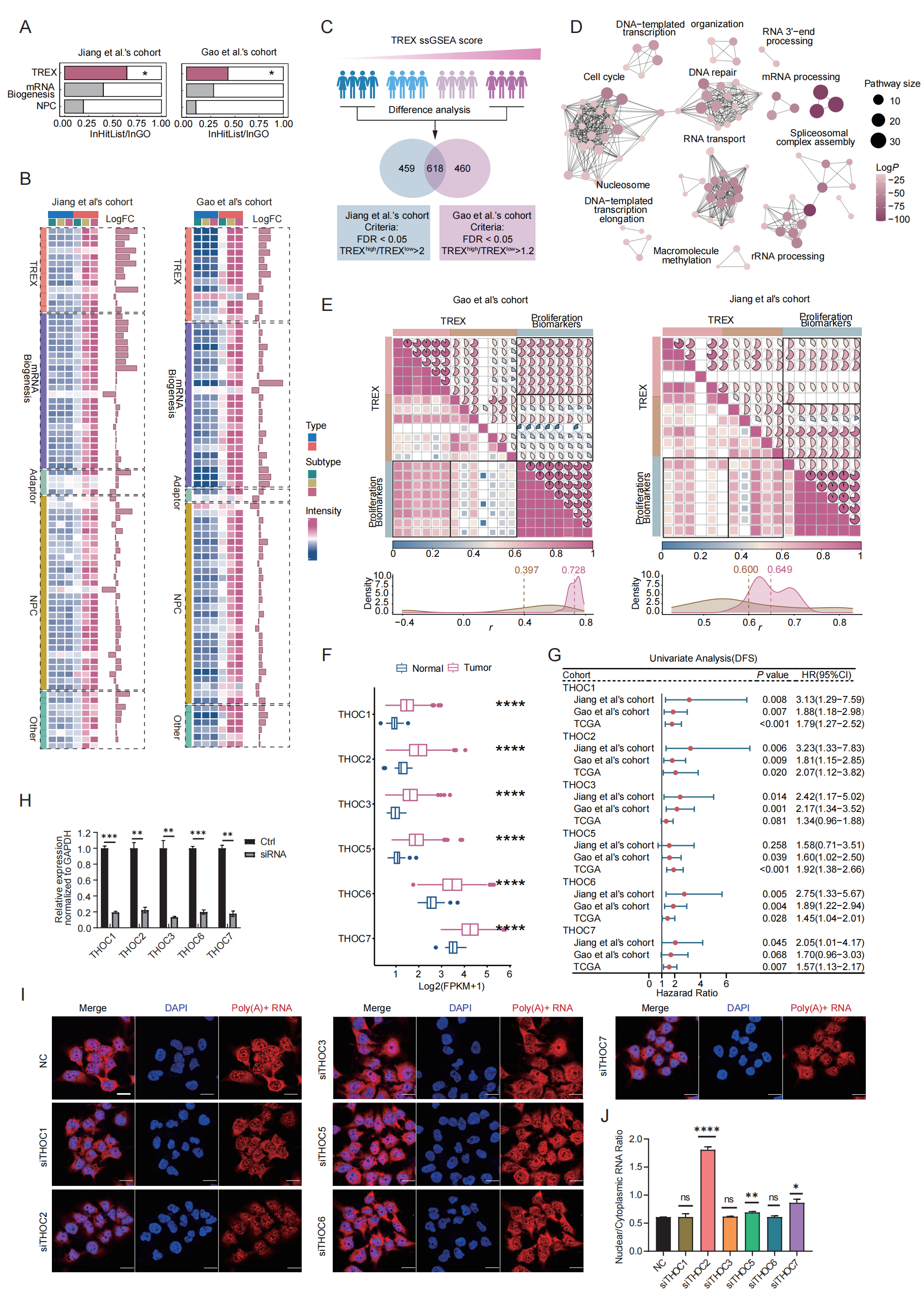

**Figure S2. THOC2 plays a critical role in regulating DNA repair in HCC, related to Fig. 2.** (**A)** mRNA biosynthesis, TREX, and NPC were used as gene sets for enrichment analysis using the results of tumor/NAT differential analysis. The horizontal axis represents the ratio of enriched proteins to the total proteins in the gene sets. (**B)** Heatmap illustrating the expression of each protein in 'Transport of Mature Transcript to Cytoplasm' in tumor and NAT samples. The histogram on the right displays the log fold change (logFC) between tumor and NAT samples. (**C)** Data analysis diagram of possible functions of TREX. The difference between the top 25% and bottom 25% of samples was analyzed based on the ssGSEA scores of TREX. The middle section featured a Venn diagram for protein screening, while the bottom part outlined the screening criteria for proteins in the two datasets. (**D)** Pathway enrichment analysis of the proteins jointly screened in (**C)**. The size of each circle represents the number of proteins contained in the enriched pathway, while the color indicates the enriched *P*-value. (**E)** The heatmap depicting the correlation between TREX proteins and proliferation markers in Gao et al.'s cohort and Jiang et al.’s cohort (above). The color and fan chart illustrate the correlation coefficient, while the labels indicate the *P*-value of the correlation. The density plot illustrates the distribution of correlation coefficients between the proliferation proteins and the two components of TREX (below), and values represent the median correlation coefficient. Rose red represents THO complex, and yellow-brown represents other parts of TREX. (**F)** Boxplot depicting the expression of each protein within the THO complex in both tumor and normal tissues, utilizing TCGA. (**G)** Forest plot of survival analysis of THO complex in two proteomics data sets and TCGA. HR > 1 indicates that a higher expression is associated with a worse prognosis. (**H)** qPCR was used to detect the knockdown effect of siRNA on each protein of the THO complex in Huh7 cells (*n* = 3). **(I)** Representative images of poly(A)+ RNA fluorescence in control (NC) and THO compex-knockdown cells detected by RNA FISH. Scale bar, 20 μm. **(J)** Quantification of nuclear/cytoplasmic RNA fluorescence intensity ratio (mean ± SEM, *n* = 3) based on RNA-FISH assays shown in **(I)**. Fisher's exact test for (**A)**; two-tailed unpaired Welch test for (**C, F, H and J)**; Spearman correlation coefficient for (**E)**; Log-rank test for (**G)**; no significance; * *P* < 0.05; ** *P* < 0.01; *** *P* < 0.001 and **** *P* < 0.0001.

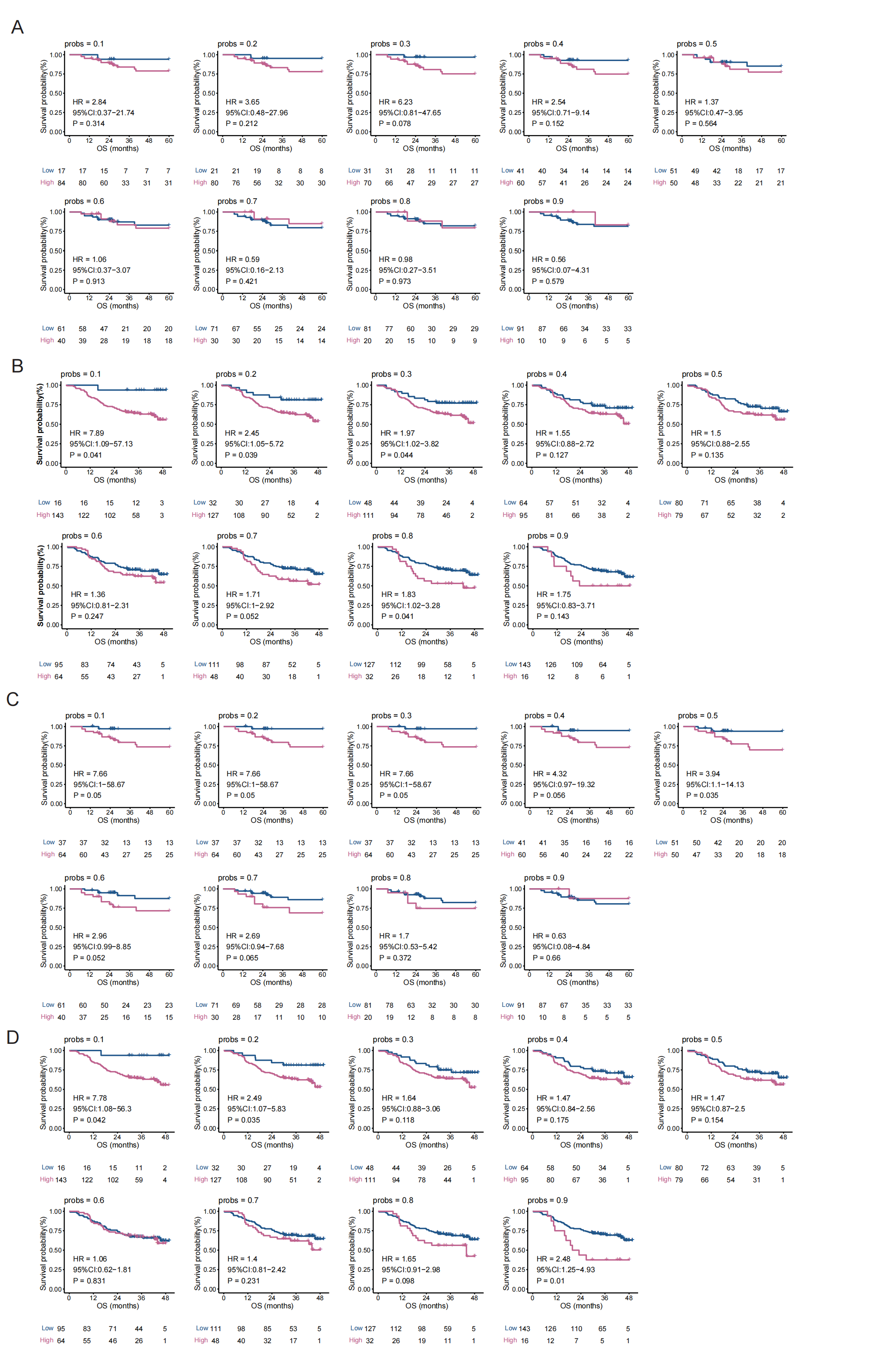

**Figure S3. Prognostic stability analyses of THOC5 and THOC7 across multiple cutoff values.** **(A-B)** Kaplan-Meier survival analyses of overall survival using multiple cutoff values (10th–90th percentile) for THOC5 expression in the Jiang et al.’s cohort **(A)** and Gao et al.’s cohort **(B)**. **(C-D)** Kaplan-Meier survival analyses of overall survival using multiple cutoff values (10th–90th percentile) for THOC7 expression in the Jiang et al.’s cohort **(C)** and Gao et al.’s cohort **(D)**. Log-rank test for **(A-D)**.

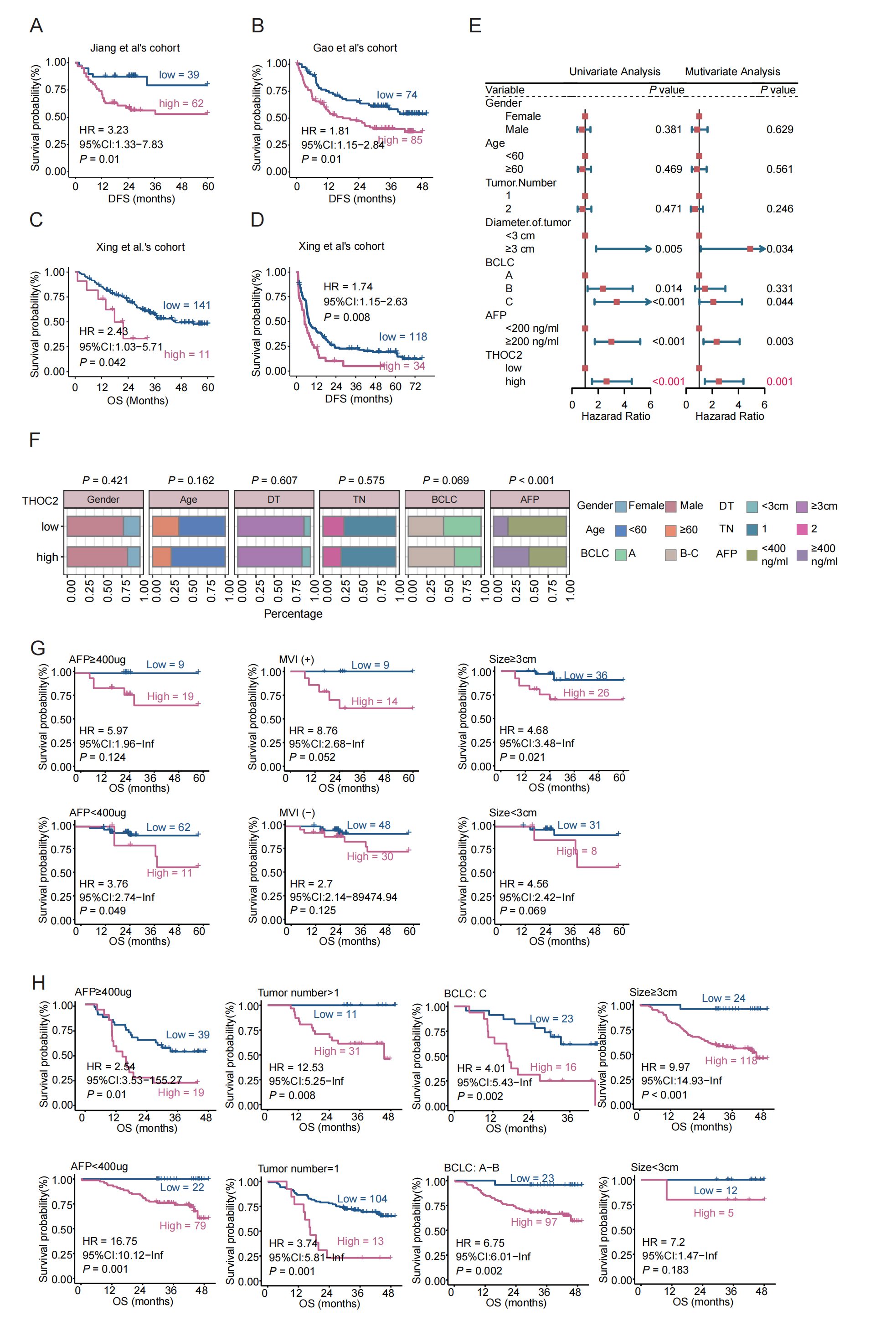

**Figure S4. THOC2 was a prognosticator for HCC patients, related to Fig. 3.** (**A, B)** Kaplan-Meier DFS curves by different THOC2 expression intensities under the optimal cut-off value in Jiang et al.’s cohort **(A)**, Gao et al.’s cohort **(B)**. (**C, D)** Kaplan-Meier OS **(C)** and DFS **(D)** curves by different THOC2 expression under the optimal cut-off value in Xing et al.'s cohort. (**E)** Univariate and multivariate Cox regression analyses of THOC2 and clinical indicators in the Gao et al.’s cohort. (**F)** The correlation between THOC2 expression and clinical indicators in Gao et al.'s cohort. Based on the cut-off value of THOC2 expression in Fig. 3B, the population was divided into high- and low-expression groups of THOC2. **(G, H)** Kaplan-Meier OS curves stratified by THOC2 expression in the Jiang et al.’s cohort **(G)** and Gao et al.’s cohort **(H)** according to clinical subgroups. DT: diameter of tumor; TN: tumor number. Log-rank test for (**A-E, G and H)**; chi-square test for (**F)**.

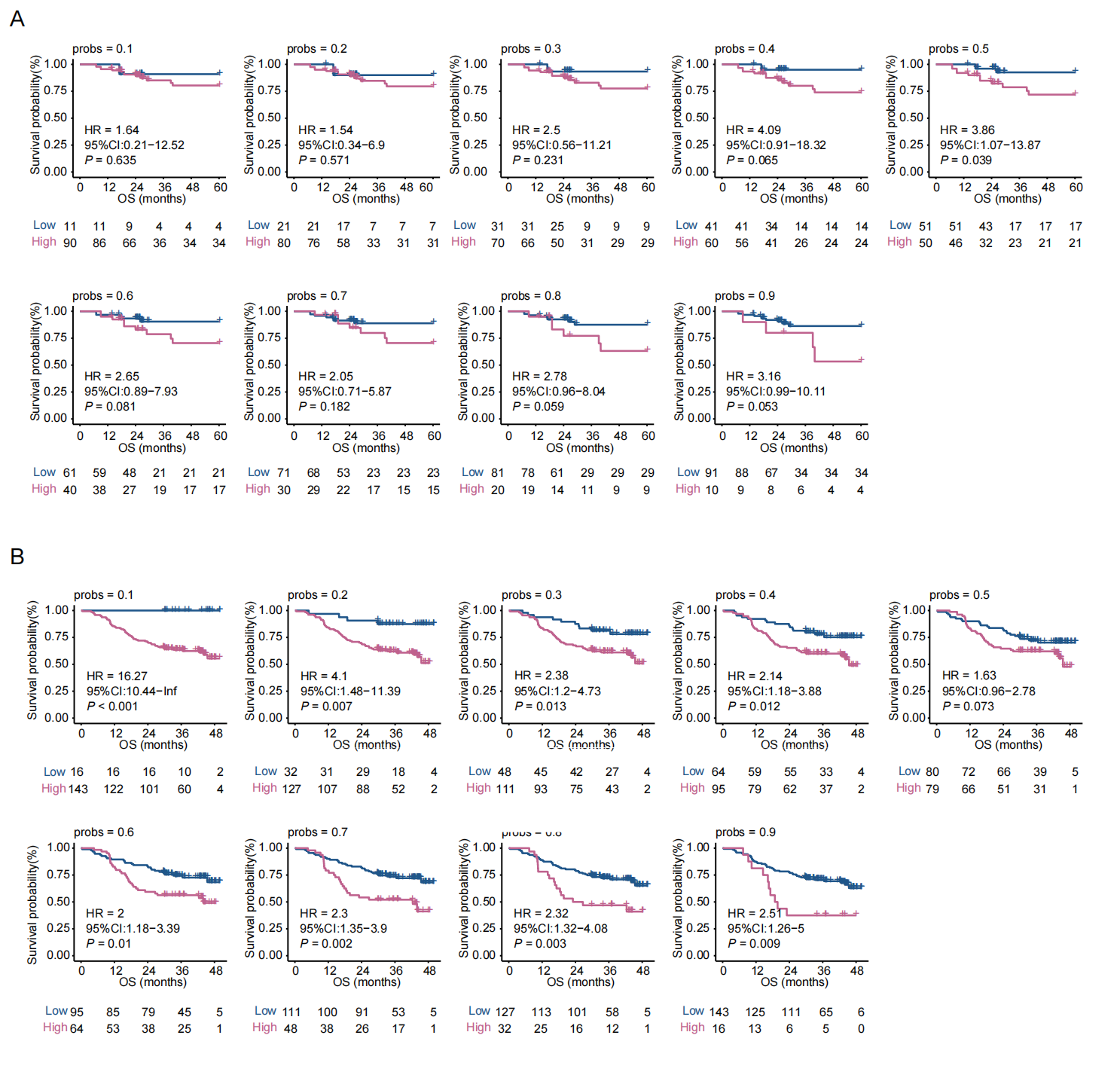

**Figure S5. Robustness of THOC2 prognostic value across multiple cutoff values.** Kaplan–Meier overall survival curves for Jiang **(A)** and Gao et al.’s cohort **(B)** stratified by THOC2 expression using a series of cutoff values ranging from the 10th to the 90th percentile (probs = 0.1–0.9). Log-rank test for **(A, B)**.

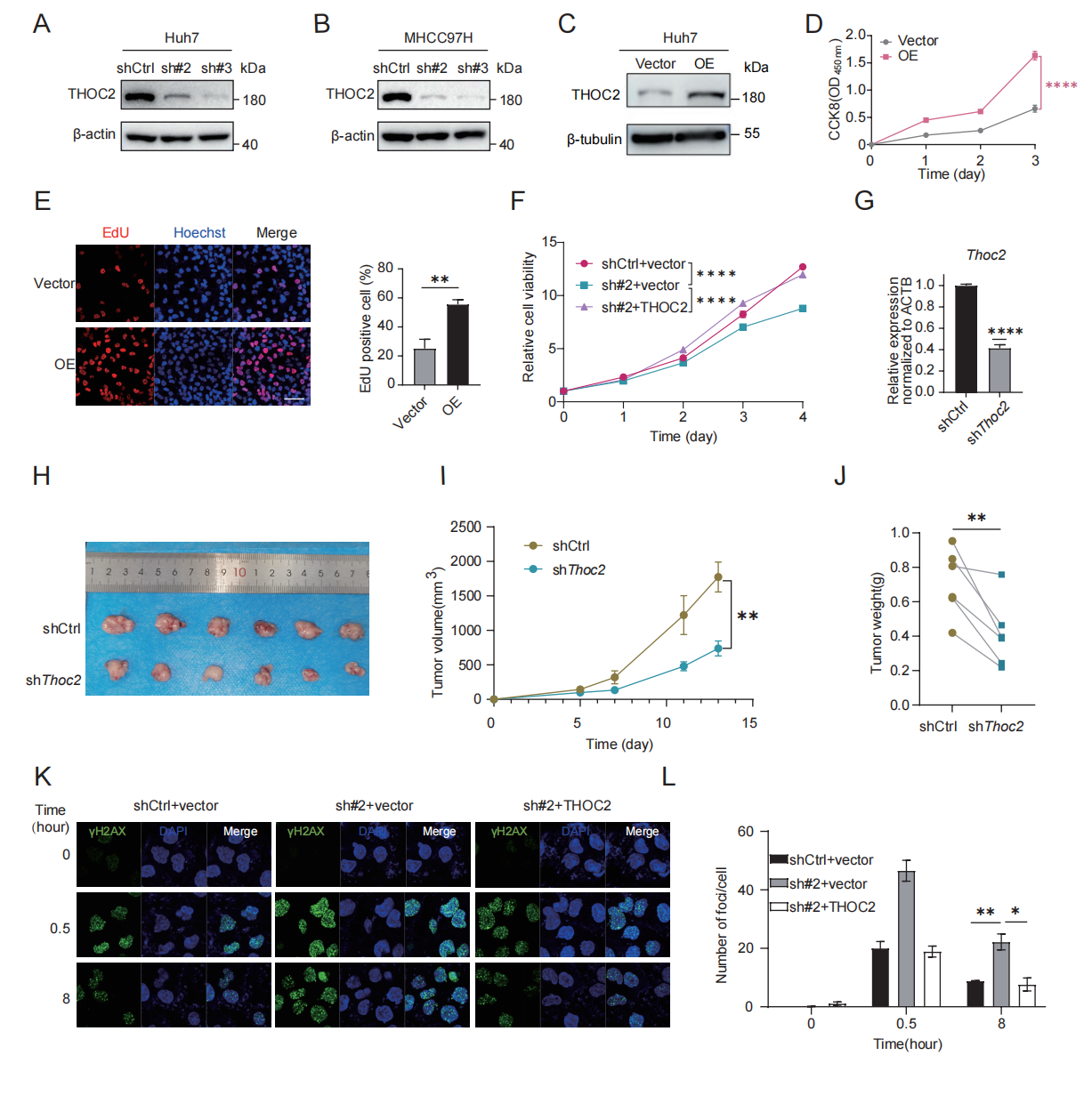

**Figure S6. THOC2 knockdown significantly inhibited tumor growth in mouse liver cancer cell, related to Fig. 4.** **(A, B)** WB analysis of THOC2 expression in THOC2-knockdown Huh7 and MHCC97H cells. **C** WB analysis of overexpressed THOC2 in Huh7 cells. (**D, E)** CCK-8 **(D)** and EdU assays **(E)** were applied to evaluate proliferation abilities of Vector and THOC2 overexpressing Huh7 cells (mean ± SEM; *n* = 3). Scale bar, 50 μm. (**F)** Cell viability was measured at the indicated time points using CCK-8 assays (mean ± SEM; *n* = 3) in control (shCtrl+vector), THOC2-knockdown (sh#2+vector), and THOC2-rescue cells (sh#2+THOC2). (**G)** qPCR detecting *Thoc2* expression in *Thoc2*-knockdown Hepa1-6 cells (mean ± SEM; *n* = 3). (**H)** Tumor xenograft models constructed from control and *Thoc2* knockdown Hepa1-6 cells (*n* = 6). (**I)** Tumor growth curve of tumor xenograft models (mean ± SEM; *n* = 6). (**J)** Tumor weight of tumor xenograft models (mean ± SEM; *n* = 6). (**K, L)** IF assay shows γH2AX foci in the control and THOC2-knockdown and THOC2-rescue cells after IR treatment (dose = 6 Gy, **K**). And histograms presented the average number of foci per cell in each group (mean ± SEM; *n* = 3, **L**). Scale bar, 20 μm. shCtrl: nontargeting shRNA; sh#2: shTHOC2-2; sh#3: shTHOC2-3; Vector: empty vector; OE: THOC2 overexpressing; two-tailed unpaired Wilcoxon t test for (**E** and **G)**; two-tailed paired Wilcoxon *t* test for (**J)**; Two way-ANOVA for (**D, F** and **I)** ; * *P* < 0.05; ** *P* < 0.01; **** *P* < 0.0001.

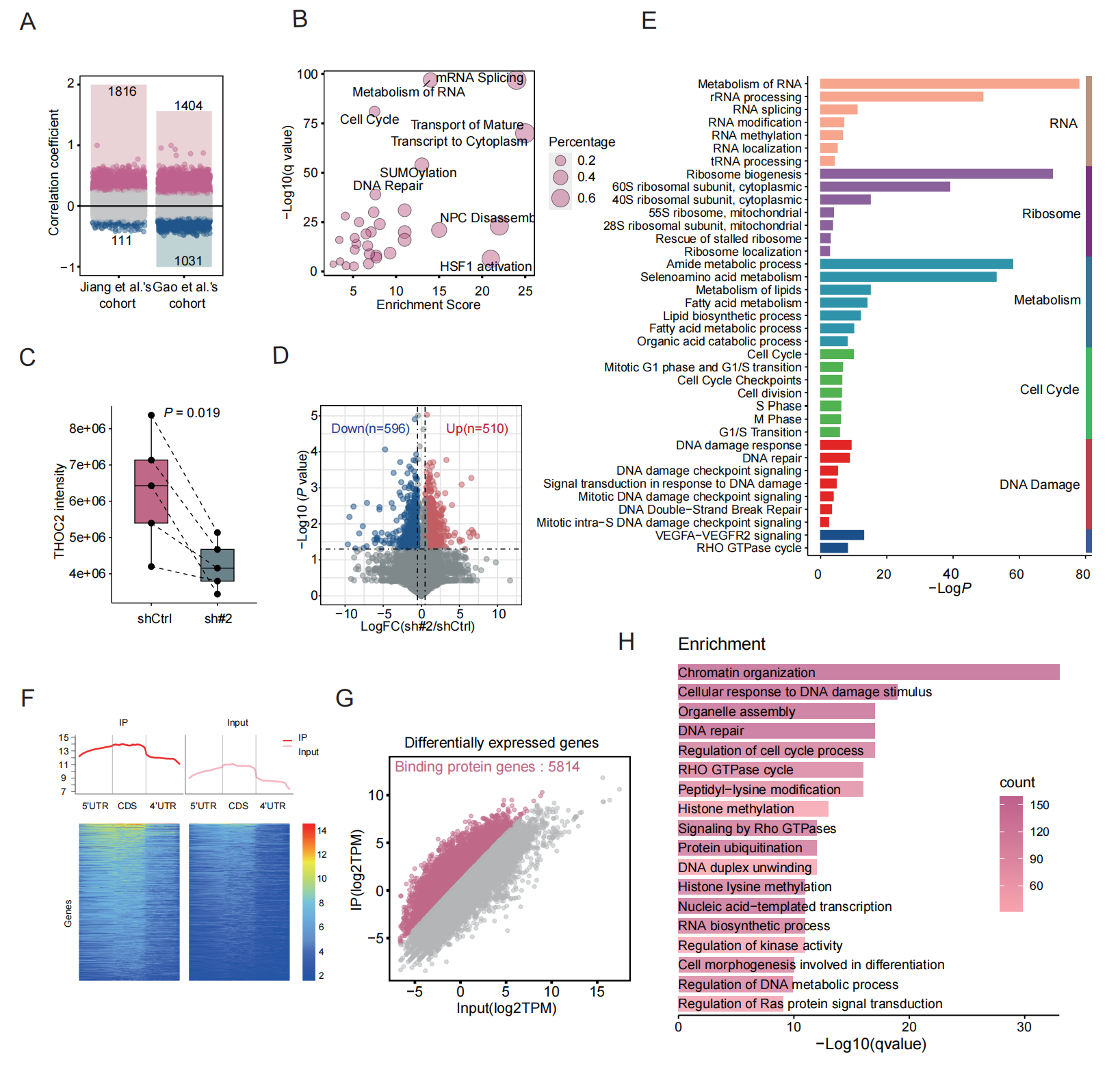

**Figure S7. Screening of mRNA types selectively exported by THOC2, related to Fig. 5.** (**A)** Co-expression proteins of THOC2 in Jiang et al.'s cohort and Gao et al.'s cohort (*P* < 0.01; *r* > 0). (**B)** Pathways enriched for co-expressed proteins in (**A)**. (**C)** THOC2 expression in subcutaneous tumors in both the control and THOC2 knockdown groups. (**D)** Volcano plot of difference analysis between control and THOC2 knockdown groups in mouse subcutaneous tumor proteomic data (*P* < 0.05; LogFC > 0.5). (**E)** Pathways enriched for up-regulated proteins in (**D)**. (**F)** Distribution of RIP-Seq reads of THOC2 in Huh7 cells. (**G)** Scatter plot of RNA expression between IP and input in RIP-seq data. Red indicates mRNAs with IP/Input > 2. (**H)** Pathways enriched in mRNAs screened by RIP-Seq data. shCtrl: nontargeting shRNA; sh#2: shTHOC2-2; Pearson correlation coefficient for (**A)**; Fisher's exact test for (**B, E** and **H)**; two-tailed paired Wilcoxon *t* test for (**C** and **D)**.

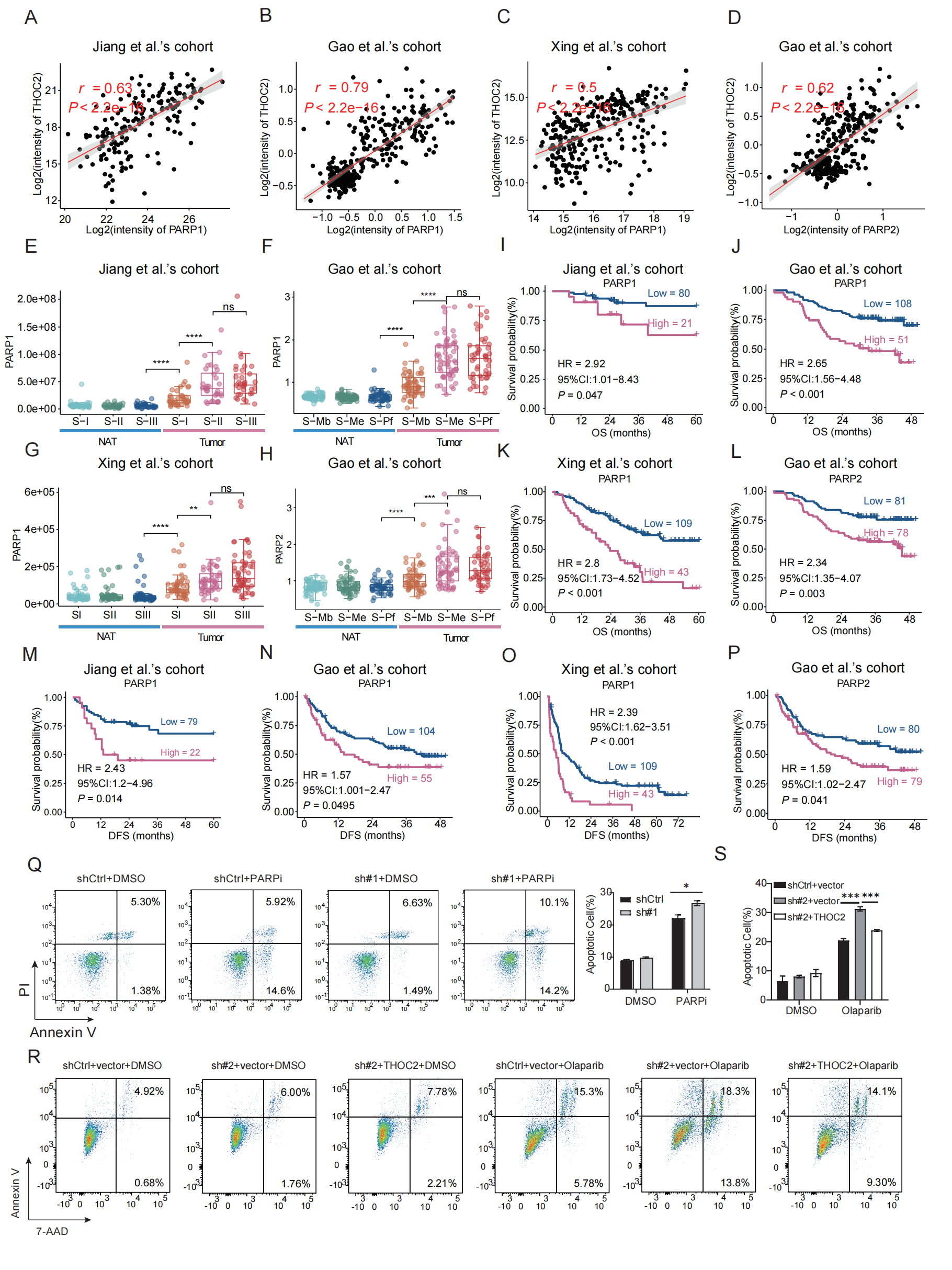

**Figure S8.** **Performance of PARP1/2 in HCC proteomics data, related to Fig. 7.** (**A-C)** Correlation between PARP1 and THOC2 expression in Jiang et al.'s cohort **(A)**, Gao et al.'s cohort **(B)** and Xing et al.'s cohort **(C)**. (**D)** Correlation between PARP2 and THOC2 expression in Gao et al.'s cohort. (**E-H)** Boxplot of PARP1/2 expression in different HCC subtypes. (**I-P)** Kaplan-Meier curve of PARP1/2 in survival analysis of three data sets. (**Q)** Apoptosis experiment after THOC2 knockdown combined with PARPi in Hepa1-6 cells. **(R, S)** Apoptosis analysis of MHCC97H cells following THOC2 knockdown and rescue under PARP inhibitor treatment (0.8 μM Olaparib). shCtrl: nontargeting shRNA; sh#2: shTHOC2-2; sh#3: shTHOC2-3; PARPi: cells treated with Olaparib (0.8 µM) which is dissolved in DMSO; DMSO: cells treated with an equal volume of DMSO to Olaparib. Pearson correlation coefficient for (**A-D)**; two-tailed unpaired Welch test for (**E-H)**; Log-rank test for (**I-P)**; two-tailed unpaired Wilcoxon *t* test for (**Q, S)**; * *P* < 0.05, ****P* < 0.001.

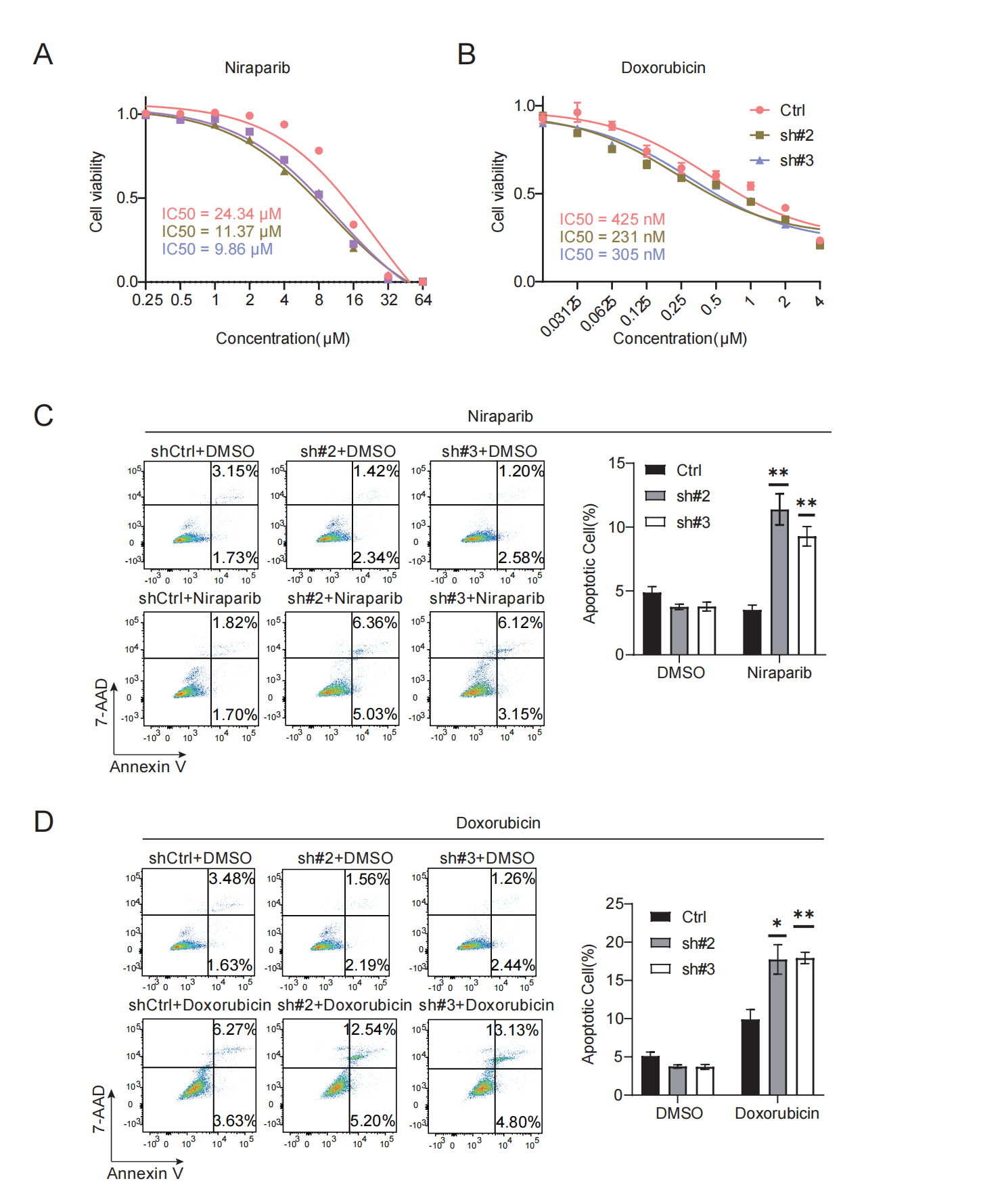

**Figure S9: THOC2 knockdown sensitizes cells to PARP inhibitors and DNA-damaging chemotherapeutic agent. (A, B)** Dose-response curves and corresponding IC50 values of Niraparib **(A)** and Doxorubicin **(B)** in control and THOC2-knockdown cells. **(C, D)** Apoptosis analysis of cells treated with Niraparib **(C)** or Doxorubicin **(D)**, as measured by flow cytometry. Ctrl, non-targeting shRNA control; shTHOC2#2 and shTHOC2#3, two independent shRNAs targeting THOC2. Data are presented as mean ± SD from at least three independent experiments. Statistical significance was determined using Student’s t-test (**P* < 0.05, ***P* < 0.01).

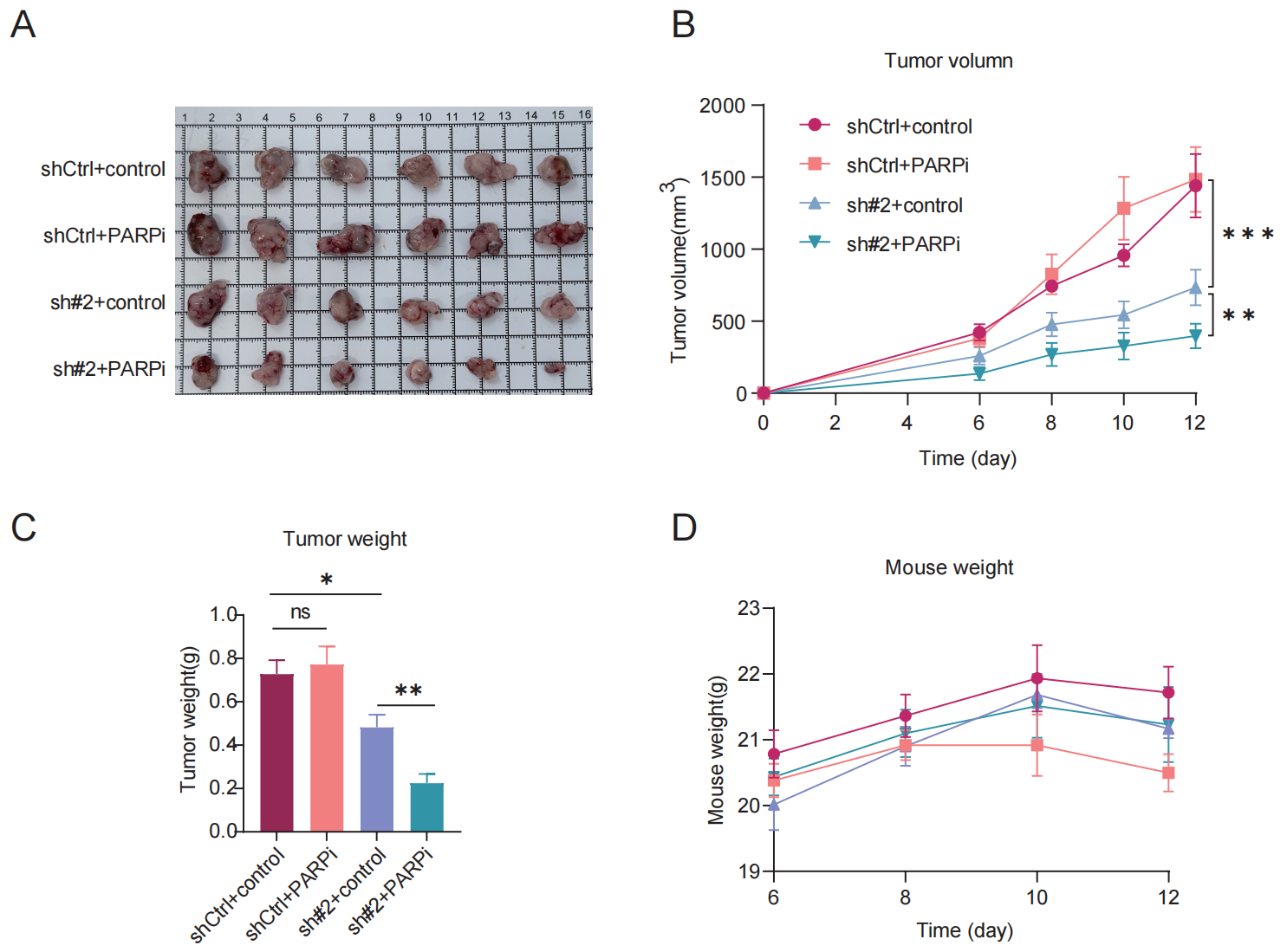

**Figure S10. Subcutaneous Hepa1-6 cells tumors (A), tumor growth curves (B) and tumor weight (C) in the indicated treatment groups (mean ± SEM; *n* = 6). (D)** Body weight changes of mice during treatment. shCtrl: nontargeting shRNA; sh#2: shThoc2-2; PARPi: mouses treated with Olaparib (50 mg/kg); Control: mouses treated with an equal volume of DMSO to Olaparib; Two-way ANOVA for **(B)**; two-tailed unpaired Wilcoxon t test for **(C)**; ns, no significance; **P* < 0.05; ** *P* < 0.01 and *** *P* < 0.001.
